# Phenotypic, molecular and functional characterisation of human *in vitro*-generated IL-17A+ CD8+ T-cells

**DOI:** 10.1101/2022.11.14.516164

**Authors:** Elizabeth H. Gray, Ushani Srenathan, Lucy E. Durham, Sylvine Lalnunhlimi, Kathryn J. A. Steel, Anca Catrina, Bruce W. Kirkham, Leonie S. Taams

**Affiliations:** Centre for Inflammation Biology and Cancer Immunology, Department of Inflammation Biology, School of Immunology & Microbial Sciences, King’s College London, London, United Kingdom; Rheumatology Unit, Department of Medicine (Solna), Karolinska Institute, Stockholm, Sweden; Department of Rheumatology, Guy’s Hospital, London, United Kingdom

**Keywords:** Tc17 cells, IL-17A, IL-17F, psoriatic arthritis, MAIT cells

## Abstract

IL-17A+ CD8+ T-cells, often referred to as Tc17 cells, have been identified at sites of inflammation in several immune-mediated inflammatory diseases including psoriasis and spondyloarthritis. Whilst much of our understanding of IL-17A+ CD8+ T-cells has been discerned from murine studies, human IL-17A+ CD8+ T-cells remain less-well characterised. We optimised an *in vitro* polarisation protocol to expand human IL-17A+ CD8+ T-cells from PBMC or bulk CD8+ T-cell populations for phenotypic and functional assessment. We show that T-cell activation in the presence of IL-1β and IL-23 significantly increased the frequencies of IL-17A+ CD8+ T-cells, which was not further enhanced by the addition of IL-6, IL-2 or anti-IFNγ mAb. *In vitro-generated* IL-17A+ CD8+ T-cells from healthy donors displayed a distinct type-17 profile compared with IL-17A- CD8+ T-cells, as defined by transcriptional signature (*IL17A, IL17F, RORC, RORA, MAF, IL23R, CCR6, CXCR6*); high surface expression of CCR6 and CD161; and polyfunctional production of IL-17A, IL-17F, IL-22, IFNγ, TNFα and GM-CSF. A significant proportion of *in vitro*-induced IL-17A+ CD8+ T-cells expressed TCRVα7.2 and bound MR1 tetramers, indicative of a MAIT CD8+ T-cell population. Using an IL-17A secretion assay, we demonstrate that the *in vitro*-generated IL-17A+ CD8+ T-cells were biologically functional and induced pro-inflammatory IL-6 and IL-8 production by synovial fibroblasts from patients with psoriatic arthritis. Collectively, we report an *in vitro* culture system to expand IL-17A+ CD8+ T-cells and further characterise their phenotype, transcriptional regulation and functional relevance to human health and disease.

## Introduction

Interleukin-17A (IL-17A) was originally identified as an effector cytokine produced by T-helper (Th) CD4+ T-cells (1) and together with fellow family member IL-17F, has come to characterise a distinct lineage termed Th17 cells. Multiple studies have described the factors and conditions that drive the differentiation of murine and human Th17 cells as well as their involvement in both host-protective and pathological immune responses (2, 3). Additionally, robust evidence implicates IL-17A (either independently or synergistically with other pro-inflammatory mediators (4, 5)) to have a pivotal function in the pathogenesis of immune-mediated inflammatory diseases such as psoriasis, psoriatic arthritis (PsA) and axial spondyloarthritis for which IL-17A is now a therapeutic target (6, 7).

*In vivo* and *in vitro* studies have highlighted IL-17A production is not restricted to Th17 cells; instead, a collection of conventional, unconventional T-cells and innate-like cell types may serve as alternative sources of IL-17A and IL-17F either in health or disease-relevant tissues (reviewed in (8, 9)). In particular, IL-17A+ CD8+ T-cells, often referred to as Tc17 cells, are known to share phenotypic and functional features of Th17 cells (reviewed in (10)). This includes expression of the lineage-committing transcription factors retinoic acid-receptor (RAR)-related orphan receptor (ROR)γt and musculoaponeurotic fibrosarcoma (c-MAF), surface expression of the typical type-17 markers CD161, CCR6 and IL-23R, and concomitant expression of Th17 (IL-17A, IL-17F, IL-21, IL-22 and granulocyte macrophage colony-stimulating factor (GM-CSF)) and cytotoxic CD8+ Tc1 (interferon (IFN)γ and tumour necrosis factor (TNF)α) related cytokines. The presence of these Tc17 cells has been described at different tissue sites in a variety of human infectious, autoimmune and inflammatory diseases, including tuberculosis (11), multiple sclerosis (12), inflammatory bowel disease (13) and psoriasis (14–16). We have also previously identified an enrichment of Tc17 cells in the joints of PsA patients that correlated with disease activity measures (17, 18). Whilst phenotypic and transcriptomic profiles of Tc17 cells derived *ex vivo* from patients with inflammatory disease have been reported, functional studies remain limited due to the relative paucity of these cells (19, 20). *In vitro* expansion of Tc17 cells could overcome this limitation and allow phenotypic and functional studies of this cell population.

To date, several murine *in vitro* studies have demonstrated that Tc17 cell differentiation can be achieved by applying Th17 polarising protocols, largely inclusive of a combination of transforming growth factor (TGF)β, IL-6, IL-1β, IL-21, IL-23, anti-IL-4 and anti-IFNγ (reviewed in (10)). Analogous to Th17 cells, murine Tc17 cells were shown to be a heterogeneous population including non-pathogenic and pathogenic subtypes, the latter typically defined by enhanced cytotoxicity with dual IL-17 and IFNγ expression promoted by IL-23 (21). However, few studies have described factors driving human Tc17 cells. The small collection of studies thus far offers a consensus in that IL-1β, IL-6, IL-23 without or with TGFβ and anti-IFNγ can direct Tc17 cell responses (11, 13, 22–26). In addition, evidence for involvement of cellular drivers comes from reports that activated monocytes from tumour sites rather than nontumour tissues more potently induced IL-17A+ CD8+ T-cells *in vitro* (27), and that pleural mesothelial cells from patients with tuberculosis infection significantly enhanced IL-17 production by patient blood-derived CD8+ T-cells, in a cell-cell contact-dependent manner (11).

Mucosal-associated invariant T (MAIT) cells are an innate-like T-cell subset defined by high expression of CD161 and their semi-invariant αβ T-cell receptor (TCR) restricted to Vα7.2-Jα33/Jα12/Jα20 (28). Expression of TCRVα7.2 restricts MAIT cells to the non-polymorphic MHC class Ib molecule MHC-related protein 1 (MR1), which presents microbial-derived metabolites of riboflavin (vitamin B2) biosynthesis including 5-(2-oxopropylideneamino)-6-ribitylaminouracil (5-OP-RU) (29). These unconventional T-cells are abundant in human tissues such as the gut and liver as well as in peripheral blood where they are predominantly CD8+ (enriched for CD8αα). Owing to their innate-like capacity, MAIT cell activation is elicited through either a TCR-dependent, TCR-independent, or synergistic manner, with the local cytokine milieu purportedly enhancing their effector function (30, 31). This innate-like functionality and particularly, a type-17 program is imparted by expression of the transcription factor promyelocytic leukemia zinc finger (PLZF) in MAIT cells as well as in other unconventional T-cells (32, 33). Subsequently, MAIT cells can express several type-17 associated markers including IL-23R, CCR6, RORγt and are potent producers of IL-17A and IL-17F that, together with IFNγ, TNFα and granzymes, rapidly orchestrate protective antimicrobial responses (reviewed in (34)). Moreover, and akin to classical Tc17 cells, MAIT cells (notably of a type-17 phenotype) have also been implicated in several inflammatory diseases including psoriasis and spondyloarthritis (35–37).

The accumulating evidence for the presence of Tc17 cells in human inflammatory diseases together with the relative paucity in knowledge regarding their induction and function provides a strong rationale for detailed characterisation of these cells. Here, we describe a protocol to polarise and increase the frequency of human IL-17A+ and IL-17F+CD8+ cells from either human PBMC or purified CD8+ T-cells, and we report their phenotypic and transcriptional profile. Furthermore, we provide evidence that human *in vitro-induced* IL-17A+ CD8+ T-cells are bioactive and elicit disease-relevant pro-inflammatory responses from PsA patient-derived synovial fibroblasts, suggestive of a functional role of these cells in PsA joint inflammation.

## Materials and Methods

### Samples and cell isolation

Human peripheral blood samples were obtained from healthy adult volunteers at King’s College London following written informed consent (Research Ethics Committee (REC) references 06/Q0705/20 and 17/LO/1940). Peripheral blood mononuclear cells (PBMC) were isolated by density gradient centrifugation using Lymphoprep™ (Axis-Shield). CD8+ T-cells were negatively isolated from PBMC by magnetic separation using the EasySep Human CD8+ T-Cell Enrichment Kit (Stemcell Technologies; average purity 93%, n=16).

Synovial fibroblast cell lines were derived from patients with PsA. Some lines were kindly provided by Professor Anca Catrina (Rheumatology Unit, Karolinska Institute, Sweden), other lines were generated in-house from PsA synovial membrane tissue obtained during knee replacement surgery at Guy’s Hospital Rheumatology Department (REC reference 07/H0809/35). In brief, to isolate synovial fibroblasts, tissue explants were cultured as 1-2mm^3^ pieces in plates coated with 0.1% bovine gelatin (Sigma-Aldrich) with DMEM (ThermoFisher Scientific) supplemented with 20% heat-inactivated fetal calf serum (FCS), 1% penicillin/streptomycin, 2% L-glutamine and 1mg/mL amphotericin B (all ThermoFisher Scientific). Tissue explants were incubated at 37°C in an atmosphere of 5% CO_2_; supplemented DMEM medium was replenished every 3 days. Synovial fibroblasts that had migrated out of the synovial tissue were collected and maintained in T175 flasks for cell line generation. Fibroblasts were passaged with1X trypsin (Sigma-Aldrich) once they reached 80% confluency and were used in cultures when between passages 3-7.

### *In vitro* type-17 polarisation

For induction of IL-17+ CD8+ T-cells, freshly isolated or cryopreserved human PBMC or purified CD8+ T-cells (1×10^6^) were cultured in complete culture medium (RPMI 1640 (Gibco) supplemented with 10% FCS, 1% penicillin/streptomycin/L-glutamine) at 37°C in an atmosphere of 5% CO_2_. Cell cultures were stimulated for 3 days with either 100 ng/mL soluble or 1.25 μg/mL immobilised (plate-bound) anti-CD3 mAb (clone OKT3, BioLegend) in combination with 1 μg/mL soluble anti-CD28 mAb (clone CD28.2, BD Biosciences) in the absence or presence of human recombinant (hr) IL-1β (10 ng/mL, Peprotech) and hrIL-23 (20 ng/mL, R&D systems). In some experiments, hrIL-6 (20 ng/mL), hrIL-2 (20U/mL, both Peprotech), neutralising anti-IFNγ or isotype control (mouse IgG2b, both 5 μg/mL, R&D Systems) were added to cultures. Cell culture supernatants were collected on day 3 before cells were restimulated with PMA/ionomycin without or with GolgiStop.

### Flow cytometric analysis

For intracellular cytokine staining, PBMC or CD8+ T-cells were stimulated, either *ex vivo* or following 3-day culture, with PMA (50 ng/mL) and ionomycin (750 ng/mL, both Sigma-Aldrich) for 3 hours at 37°C in the presence of GolgiStop (monensin, according to manufacturer’s recommendation, BD Biosciences). Cells were washed and labelled with fixable viability dye (LIVE/DEAD eFlour780, eBioscience) for 15 min at 4°C. PBMC samples were then FcR-blocked using 10% human AB serum (Invitrogen) in FACS buffer for 15 min at room temperature (RT). Where applicable, cells were stained with human MR1-5-OP-RU or MR1-6-FP (supplied by the NIH Tetramer Core Facility) for 40 min at RT followed by incubation at 4°C for 30 min with a combination of mAbs against cell surface markers including: CD8, CD14, CD19, CD161, CCR6, and TCRVα7.2 (full details described in **Supplementary Table 1**). Cells were fixed with 2% paraformaldehyde for 15 min at RT then permeabilised using 0.5% saponin (Sigma-Aldrich) and stained for 30 min at 4°C for the following intracellular markers: CD3, CD4, IL-17A, IL-17F, IFNγ, TNFα, GM-CSF, granzyme A and B (full details in **Supplementary Table 1**). Samples were acquired using either a FACS Canto II or LSRFortessa (BD Biosciences) and data were analysed using FlowJo software (v10.7.1, TreeStar Inc.). CD8+ T-cells were gated as shown in **Supplementary Figure 1**. FM control stainings were used to aid determination of cytokine-expressing cell populations. Boolean gating strategy of IFNγ, TNFα and GM-CSF expression by IL-17A+ Vα7.2-CD8+ and IL-17A+ Vα7.2+ CD8+ T-cell subsets was imported into SPICE software (v5.1) for visualisation of cells expressing polyfunctional or monofunctional cytokine combinations.

### IL-17A cytokine secretion assay (CSA)

For transcriptional and functional assessments, IL-17A-producing CD8+ T-cells were isolated post-polarising culture using an IL-17A cytokine secretion assay (Miltenyi Biotech, according to manufacturer’s recommendations) combined with FACS sorting. Purified CD8+ T-cells were cultured for 3 days with plate-bound anti-CD3 and soluble anti-CD28 mAbs in the presence of hrIL-1β and hrIL-23. After 3 days, cells were re-stimulated with PMA (50ng/mL) and ionomycin (750ng/mL) for 1.5 hours before cells were counted with trypan blue. Cells were washed then resuspended in cold complete culture medium and labelled with the IL-17A catch reagent for 5 min on ice. Cell suspension was diluted with warm complete culture medium and incubated in a continuous motion using the MACSmix rotator (Miltenyi Biotec) for 45 min at 37°C (5% CO_2_) to allow secretion and capture of IL-17A on the cell surface-bound catch antibody. After cells were washed with cold PBS containing 0.5% EDTA, they were labelled for viability, CD3, CD4, CD8 (with CD19 as an exclusion marker) as well as the IL-17A detection antibody (PE or APC conjugated) for 20 min on ice. Once washed, IL-17A-secreting (IL-17A+) and non-IL-17A-secreting (IL-17A-) CD8+ T-cells were immediately FACS sorted on a BD FACS ARIA II. In some experiments, mAbs against the semi-invariant TCR Vα7.2 (with pan γδTCR and CD56 as exclusion markers) were included in the panel to sort IL-17A+ Vα7.2- and IL-17A+ Vα7.2+ cells from CD8+ T-cell cultures. Sorted T-cell subsets were either stored in TRIzol®Reagent (ThermoFisher) at −80°C for later qPCR array analysis or used directly for functional assessments. For the latter, sorted T-cells were either added to fibroblasts as described below or cultured for 20 hours at 37°C (5% CO_2_) in complete culture medium for supernatant generation.

### PsA synovial fibroblast co-cultures

PsA synovial fibroblasts were seeded (1×10^4^ per well) in a flat-bottomed 96-well plate in supplemented DMEM medium and incubated for 24 hours at 37°C (5% CO_2_). Following supernatant removal, fibroblasts were cultured in supplemented DMEM in the absence or presence of 20% (v/v) cell culture supernatants from FACS-sorted CD8+ T-cell populations for a further 24 hours, after which supernatants were collected and analysed for IL-6 and IL-8 production. Alternatively, fibroblasts were co-cultured for 24 hours with CSA-FACS sorted IL-17A+ or IL-17A- CD8+ T-cells at a 1:2.5 fibroblast to T-cell ratio, in the absence or presence of anti-IL-17A mAb (secukinumab, Novartis) and/or anti-TNFα blocking antibodies (adalimumab, Abbott Laboratories) or isotype control mAb (all human IgG1 and added at 5μg/mL).

### Cytokine detection

The presence of IL-17A in T-cell culture supernatants and of IL-6 and IL-8 in fibroblast culture supernatants was quantified by ELISA according to manufacturer instructions (deluxe or standard kits, respectively, all BioLegend). Plates were read at 450 nm using a Spark 10M (Tecan). Cytokine secretion profiles of supernatants from FACS-sorted IL-17A+ Vα7.2- CD8+, IL-17A+ Vα7.2+ CD8+ and IL-17A- CD8+ T-cell subsets were assessed by custom magnetic Luminex (Bio-Techne). Luminex plates were analysed on a Luminex FlexMap 3D platform. IL-17A and IL-17F were measured on separate assay plates due to the cross-reactivity of the magnetic beads with IL-17AF.

### RNA extraction and cDNA synthesis

Total RNA was extracted from healthy donor CSA-FACS sorted IL-17A+ CD8+ and IL-17A- CD8+ T-cell subsets (4,175 – 43,856 cells or 1×10^6^ cells, respectively) using the TRIzol®Reagent phenol-chloroform extraction method in combination with Phasemaker tubes (ThermoFisher). To increase total RNA precipitation, during the isopropanol step, samples were kept at −80°C for 24 hours and 1 μl GlycoBlue^TM^ Coprecipitant (ThermoFisher) was added. RNA yield and integrity (mean RIN = 8.9) were assessed by Bioanalyzer (Waterloo Genomics Centre, KCL) and RNA was stored short-term at −80°C. Amount of RNA input for complementary DNA (cDNA) transcription was standardised within donor paired samples by adjusting RNA concentrations to the total extracted from the IL-17A+ CD8+ T-cell subset (range 30.2ng – 213.0ng). First-strand cDNA synthesis was performed as a 20 μl reaction using the LunaScript^®^ RT SuperMix Kit (NEB, following the manufacturer’s protocol) and cDNA was stored short-term at −20°C before transcriptional assessment.

### Quantitative real-time polymerase chain reaction (RT-qPCR)

Gene expression profiles of *in vitro*-generated IL-17A+ CD8+ and IL-17A- CD8+ T-cells were assessed by bespoke TaqMan^®^ qPCR array (all 96 primer assays including 3 endogenous controls and 93 targets, were selected from ThermoFisher, details listed in **Supplementary Table 2**). Array cards and sample cDNA templates were prepared according to manufacturer’s instructions using the TaqMan^®^ Fast Advanced PCR Master Mix (ThermoFisher). RT-qPCR was performed using a QuantStudio^TM^ 7 Flex System (Applied Biosystems) with the following amplification conditions: hold at 50°C for 2 min then hold at 92°C for 10 min followed by 40 cycles of 95°C for 1 sec and 60°C for 20 sec. Housekeeping genes 18S ribosomal RNA (*18S*), beta-2-microglobulin (*B2M*) and peptidylprolyl isomerase A (*PPIA*) were screened for stability using the web-based tool RefFinder and R-based package NormFinder. Expression of each target gene was normalised to the expression of *B2M* and *PPIA* within a sample, performed using the comparative threshold cycle (Ct) method calculated as: ΔCt = Ct(gene of interest) - Ct(geometric mean *B2M* + *PPIA*). Ct was defined as 40 for the ΔCT calculation when the signal was under detectable limits. The relative fold change in target mRNA expression levels was assessed in IL-17A+ CD8+ versus IL-17A- CD8+ T-cells and calculated with the formula 2^-ΔΔCT^ where for a given gene ΔΔCT =ΔCt(IL- 17A+) –ΔCt(IL-17A-).

### Data analysis and statistical testing

Graphs were constructed and statistical tests performed using GraphPad Prism v9.1. Sample sizes with n<8 or when the population assumed a non-normal distribution were tested non-parametrically using a Wilcoxon signed-rank matched pairs test. Data sets with n>8 were tested for normality using the D’Agostino and Pearson omnibus normality test then tested for significance using the appropriate parametric or non-parametric test as stated in the figure legends. For transcriptional data analysis, undetectable genes (*ZBTB32* and *CD5L*) were filtered from initial principal-component analysis and heatmap generation; both were computed in R using prcomp and pheatmap functions. Genes that were identified to have very low/negligible expression in both IL-17A+ CD8+ and IL-17A- CD8+ T-cells with reference to the geometric mean of *CD4* normalised expression, were excluded from statistical testing and relative fold change analysis (*IL17B, IL17C, IL17D, IL25, IL17RC* and *CCR9*). Differential statistical analysis was performed on normalised expression values using a parametric paired Student’s t-test with Holm-Šídák multiple comparisons test with adjusted p-values (p<0.05) reported.

## Results

### Human peripheral blood IL-17+ CD8+ T-cells are detected at low frequencies *ex vivo* and can be expanded *in vitro* in the presence of IL-1β and IL-23

We first sought to determine the *ex vivo* frequencies of IL-17A+ and IL-17F+ CD8+ T-cells in human peripheral blood. Freshly isolated or cryopreserved healthy donor PBMC were stimulated for 3 hours with PMA/ionomycin in the presence of GolgiStop followed by intracellular cytokine staining. CD8+ T-cells were gated as shown in **Supplementary Figure 1A**. Low frequencies of IL-17A+ cells (median 0.08%), and very low frequencies of IL-17F+ cells (median 0.007%), were detected within the CD8+ T-cell population *ex vivo* (**Figure 1A, B**). In contrast, higher frequencies of IL-17A+ and IL-17F+ cells were detected in peripheral blood CD4+ T-cells (median 0.75% and 0.04%, respectively, **Supplementary Figure 2A, B**).

**Figure 1.**
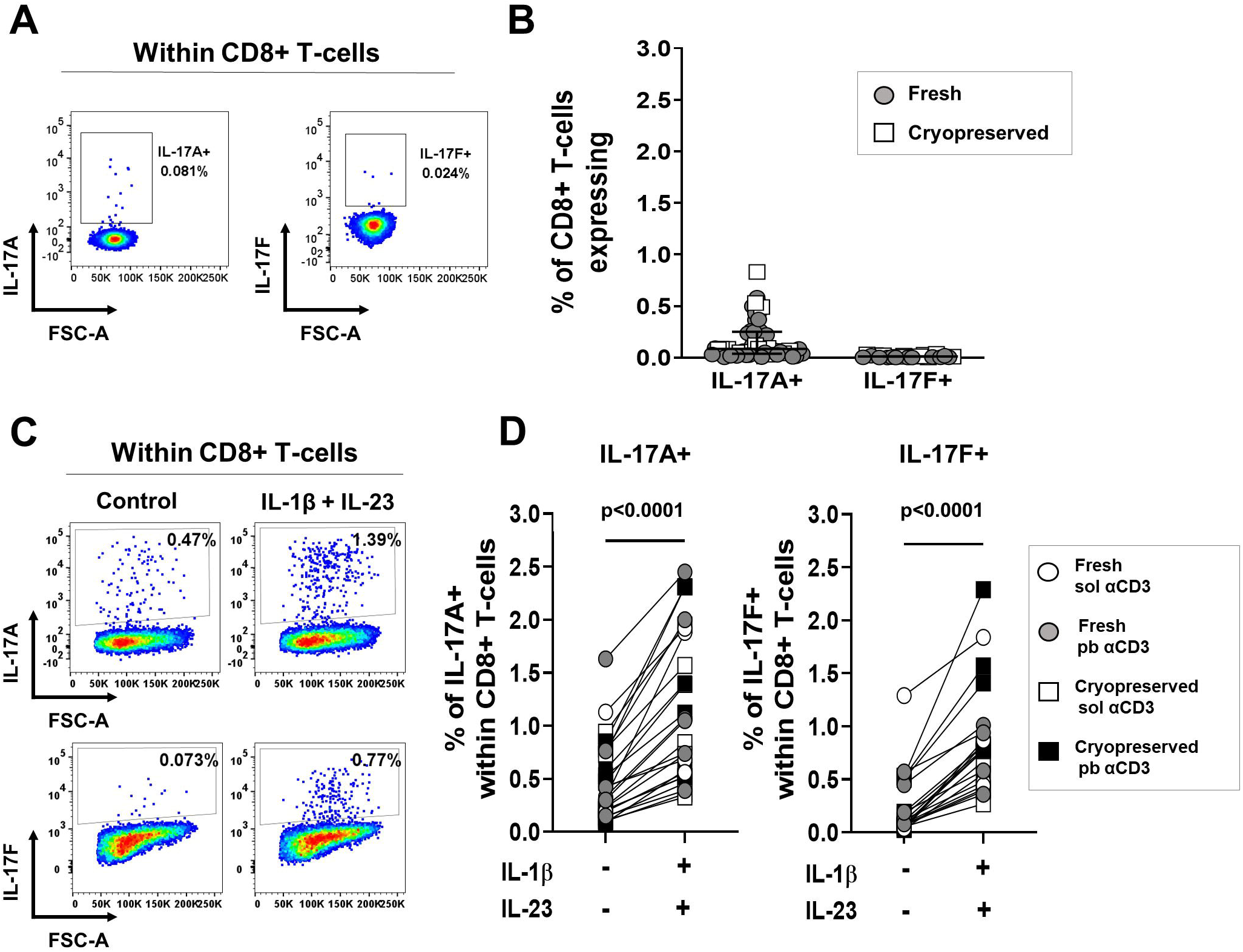
Low *ex vivo* frequencies of IL-17A+ and IL-17F+ CD8+ T-cells in healthy donor PBMC are expanded upon anti-CD3/CD28 stimulation in the presence of IL-1β and IL- 23. **(A, B)** Freshly isolated (circles) or cryopreserved (squares) healthy donor PBMC were stimulated *ex vivo* for 3 hours with PMA/ionomycin and GolgiStop for assessment of intracellular IL-17A and IL-17F cytokine expression by flow cytometry. Representative staining plots **(A)** and cumulative data **(B)** show frequencies of IL-17A+ and IL-17F+ cells within live CD3+CD8+ T-cells from independent donors (n=51 and n=23, respectively). **(C, D)** Freshly isolated (circles) or cryopreserved (squares) healthy donor PBMC were cultured for 3 days with either plate-bound (filled symbols) or soluble (open symbols) anti-CD3 mAb and soluble anti-CD28 mAb in the absence (control) or presence of hrIL-1β (10 ng/mL) and hrIL- 23 (20 ng/mL). After 3 days, cells were re-stimulated with PMA/ionomycin and GolgiStop for detection of intracellular cytokine expression. Representative dot plots **(C)** and cumulative data **(D)** show the frequencies of *in vitro*-induced IL-17A+ and IL-17F+ cells within live CD3+ CD8+ T-cells from independent donors (n=50 and n=22, respectively). Statistical analysis performed using Wilcoxon matched-pairs signed rank test.

To investigate whether IL-17+ CD8+ T-cells could be expanded *in vitro*, freshly isolated or cryopreserved PBMC from healthy donors were stimulated using plate-bound or soluble anti- CD3 mAb with soluble anti-CD28 mAb in the absence or presence of the well-established human Th17-promoting cytokines IL-1β and IL-23. After 3 days, cells were re-stimulated with PMA/ionomycin and Golgistop for intracellular cytokine assessment by flow cytometry (CD8+ T-cell gating strategy shown in **Supplementary Figure 1B**). Frequencies of IL-17A+ and IL- 17F+ cells were significantly increased when PBMC were cultured in the presence of IL-1β and IL-23, as compared with cell cultures with only anti-CD3/CD28 stimulation (3.2-fold and 7-fold higher respectively, both p<0.0001) (**Figure 1C, D**). As expected, culturing PBMC under type-17 polarising conditions led to a significant increase in IL-17A+ and IL-17F+ CD4+ T-cells also (p<0.0001, **Supplementary Figure 2C, D**).

### *In vitro* induction of IL-17+ CD8+ T-cells upon anti-CD3/CD28 stimulation in the presence of IL-1β and IL-23 is not further enhanced by addition of IL-6, IL-2 or anti-IFNγ

Given that the presence of other immune cell subsets in whole PBMC cultures may have contributed additional type-17 promoting and/or inhibitory factors, we next sought to directly explore the effect of IL-1β and IL-23 upon IL-17A+ CD8+ T-cell induction. For this, CD8+ T-cells were purified by magnetic bead separation from freshly isolated healthy donor PBMC and cultured for 3-days with plate-bound anti-CD3 and soluble anti-CD28 mAbs in the absence or presence of IL-1β and IL-23 (purity assessment and representative gating strategy shown in **Supplementary Figure 1C, D**). Akin to whole PBMC cultures, a statistically significant increase in IL-17A+ cells was observed when CD8+ T-cells were stimulated in the presence of type-17 polarising cytokines as opposed to anti-CD3/CD28 stimulation alone (median 1.44% versus 0.28%, 5.1-fold increase, p=0.0005,) (**Figure 2A**). This was confirmed at the cytokine secretion level, with significantly elevated levels of IL-17A detected in cell culture supernatants of CD8+ T-cells cultured under type-17 polarising conditions (469 pg/mL versus 99 pg/mL, 4.7-fold increase, p=0.0313) (**Figure 2B**).

**Figure 2.**
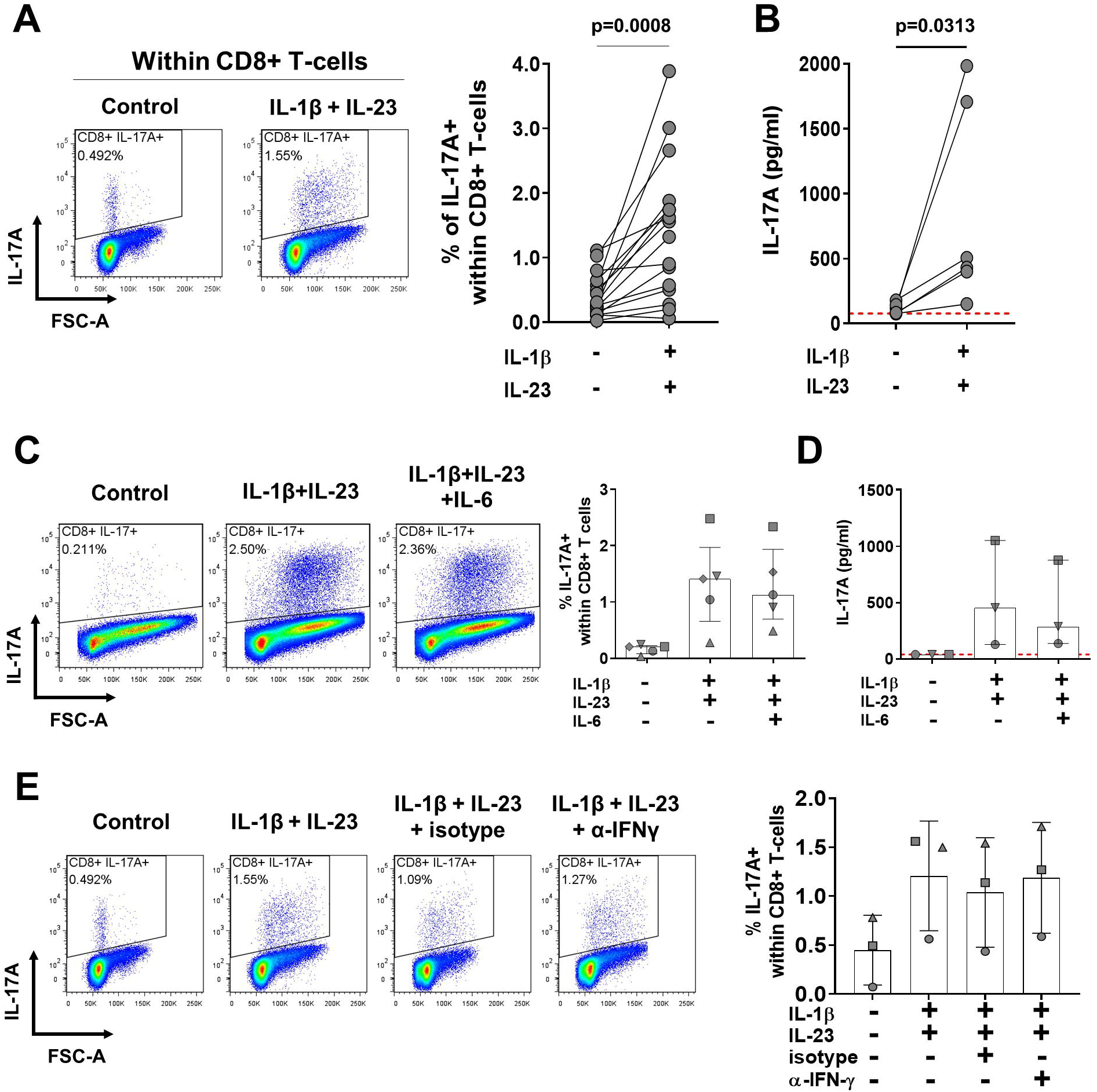
Anti-CD3/CD28 stimulation in the presence of IL-1β and IL-23 is sufficient to expand IL-17A+ CD8+ T-cells from purified CD8+ T-cell cultures. Purified CD8+ T-cells from freshly isolated PBMC were cultured in the presence of plate-bound anti-CD3 mAb and soluble anti-CD28 mAb in the absence (control) or presence of hrIL-1β and hrIL-23 for 3 days. On day 3, culture supernatants were removed and remaining cells were re-stimulated for 3 hours with PMA/ionomycin and GolgiStop for detection of intracellular IL-17A expression by flow cytometry. **(A)** Representative staining plots and cumulative data show frequencies of IL- 17A+ cells within live CD3+ CD8+ T-cells (n=16). **(B)** Levels of IL-17A detected in 3-day culture supernatants were measured by ELISA (n=6). Statistical analysis was performed using a paired t-test **(A)** or Wilcoxon matched-pairs signed rank test **(B)**. **(C-E)** Bulk CD8+ T-cells were cultured under type-17 polarising conditions without or with hrIL-6 (20 ng/mL) **(C, D)** or in the absence or presence of either neutralising anti-IFNγ or isotype control mAb (5 μg/mL) **(E)** for 3 days. Representative stainings and cumulative data (n=3-5, each symbol correspond to an independent donor) show frequencies of IL-17A+ cells within live CD3+ CD8+ T-cells as detected by flow cytometry **(C and E)**; or **(D)** indicate levels of IL-17A detected by ELISA in culture supernatants collected prior to re-stimulation (n=3). Data are plotted as median + IQR.

Previously, human and murine studies have shown that IL-6 can promote Th17 and IL-17A+ CD8+ T-cell polarisation, whilst IFNγ can antagonise this (38, 39). However, the addition of hrIL-6 or of IFNγ blocking antibodies to our purified CD8+ T-cell culture system did not further increase the frequency of IL-17A+ cells or the production of IL-17A by CD8+ T-cells (**Figure 2C-E**). Addition of hrIL-2 and extending the culture system to 6 days also did not achieve any further consistent additive effect on the frequency of induced IL-17A+ CD8+ T-cells (**Supplementary Figure 3**).

### *In vitro*-induced IL-17A+ CD8+ T-cells are characterised by a core type-17 transcriptional and phenotypic signature

We next sought to determine whether human *in vitro*-generated IL-17A+ CD8+ T-cells display a type-17-related gene profile compared with IL-17A- CD8+ T-cells. For this, after polarising culture, highly pure IL-17A-secreting (IL-17A+) and non-IL-17A-secreting (IL-17A-) CD8+ T-cells were FACS sorted using an IL-17A cytokine secretion assay (CSA) (representative gating strategy shown in **Supplementary Figure 4A**). Accuracy of the combined CSA-FACS sorting method was validated by assessment of secreted IL-17A in cell culture supernatants from sorted IL-17A+ and IL-17A- CD8+ T-cell subsets, as well as by determining expression of *CD8A, CD8B* and *CD4* for each population (**Supplementary Figure 4B, C**).

We investigated the expression of a broad range of type-17 associated genes using a bespoke qPCR array. First, we applied principal-component analysis (PCA) to globally evaluate transcriptomic profiles of the sorted *in vitro*-induced T-cell subsets. IL-17A+ CD8+ T-cells from all 7 independent donors clustered separately from IL-17A- CD8+ T-cells (upper panel **Figure 3A**). Several hallmark type-17 genes were identified among the top 20 genes that contributed to component 1, which accounted for most of the variation described by the PCA plot (PC1, 47.3%) (lower panel **Figure 3A**). Heatmap analysis further highlighted that a type-17-related gene signature was enriched in all donor IL-17A+ versus IL-17A- CD8+ T-cells (**Figure 3B**). Quantitative assessment deduced that a total of 29/85 genes were statistically more abundant in *in vitro*-polarised IL-17A+ versus IL-17A- CD8+ T-cells (select genes shown in **Figure 3C,** all 29 listed in **Supplementary Table 3**). Importantly, a large proportion of these enriched genes are characteristic of the type-17 program including: cytokines and chemokines *IL17A, IL17F, IL26 and CCL20;* migratory, lineage-defining and signalling receptors *CCR6, CXCR6, KLRB1* (encoding CD161) *and IL23R;* as well as transcription factors *RORC, RORA* and *MAF* (encoding RORγt, RORα and c-MAF, respectively). We also detected slightly increased expression of *IL2RA, CTLA4, PDCD1* (encoding programmed cell death 1 (PD-1)) and *GZMB* in polarised IL-17A+ CD8+ T-cells versus IL-17A- counterparts. Our analysis did not identify significant differences in mRNA expression of additional type-17 effector molecules *IL21, IL22 and CSF2* (encoding GM-CSF) or in *IFNG* or *TNF* which both displayed high mRNA levels in each T-cell subset. Transcript levels of *TCF7* (encoding T-cell factor 1 (TCF-1) a recently identified murine Tc17 cell transcriptional regulator (40)) was variable among donor IL-17A+ CD8+ T-cells and not significantly lower than levels detected in IL-17A- CD8+ T-cells. Evaluation of other T-cell lineage specific markers namely *TBX21* (encoding T-bet)*, CXCR3, GATA3, FOXP3* and *IL10* showed no selective expression in either subset.

**Figure 3.**
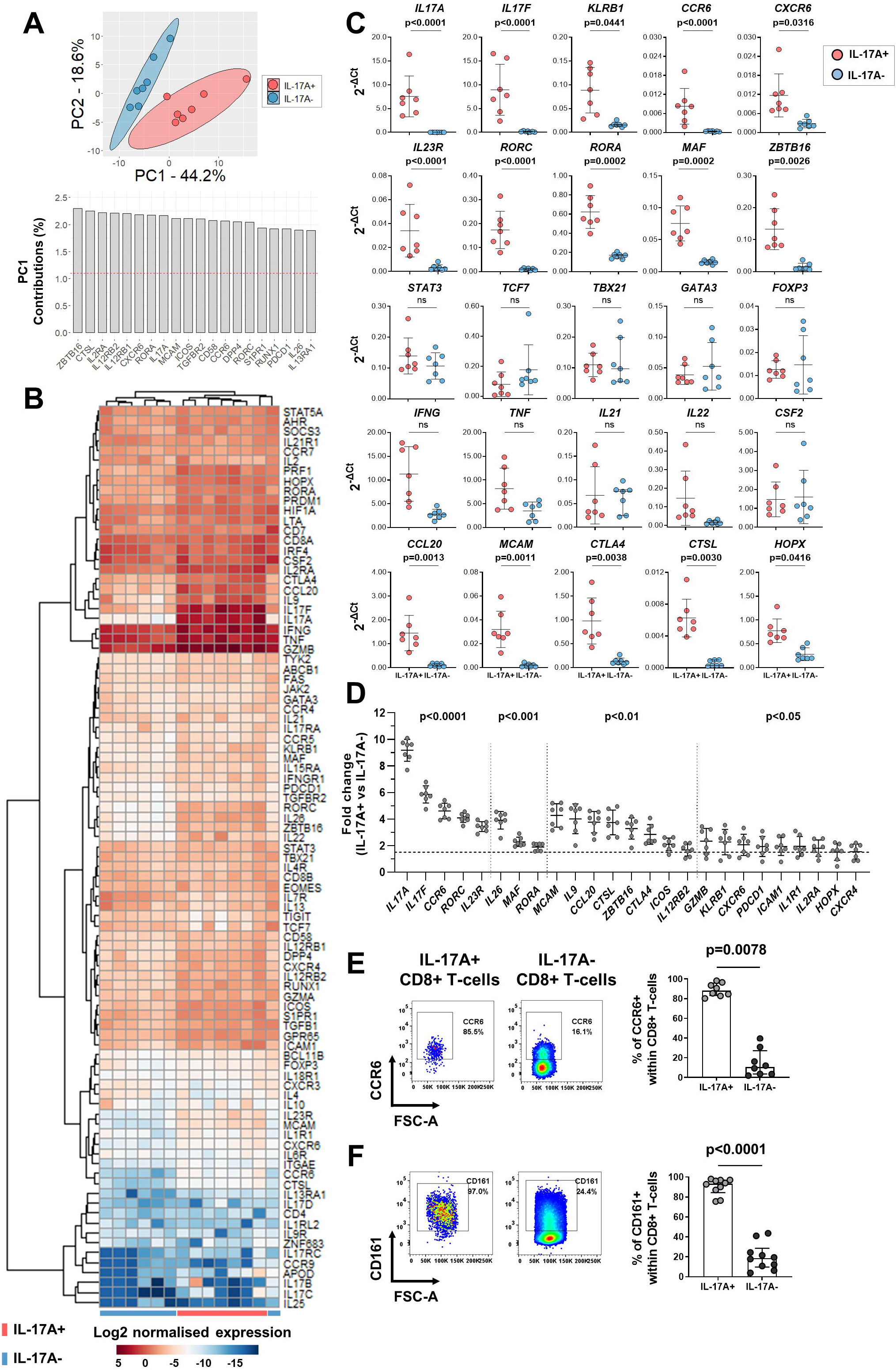
Transcriptional analysis reveals *in vitro*-generated IL-17A+ CD8+ T-cells acquire a signature type-17-related profile. Bulk CD8+ T-cells were isolated from healthy donor PBMC and cultured for 3 days with plate-bound anti-CD3 and soluble anti-CD28 mAbs in the presence of hrIL-1β and hrIL-23. On day 3, cells were re-stimulated for 1.5 hours with PMA/ionomycin for IL-17A CSA combined FACS sorting of *in vitro*-generated IL-17A+ CD8+ and IL-17A- CD8+ T-cells. **(A-D)** Gene profiles of matched donor T-cell subsets were assessed by custom qPCR array (n=7 independent donors). Each target gene expression was normalised to endogenous controls *B2M* and *PPIA* **(A)** Principal-component analysis (PCA) of IL-17A+ (red) and IL-17A- (blue) CD8+ T-cell transcriptional profiles plotted on components 1 and 2 (upper panel); bar plot of the top 20 PCA parameter contributions (%) for component 1 (lower panel). **(B)** Heatmap showing gene expression profiles in IL-17A+ compared with IL-17A- CD8+ T-cells with donors assigned columns and rows the log2 transformed normalised gene expression. Unbiased hierarchical gene and sample clustering indicated by dendrograms. **(C)** Scatter plots depict absolute mRNA expression levels (2^-ΔCt^) of a selection of T-cell lineage specific genes in IL-17A+ (red) and IL-17A- (blue) CD8+ T-cells. **(D)** Scatter plot shows relative fold change (2^-ΔΔCt^) in type-17 signature genes identified as significantly differentially expressed with fold change > 1.5 (marked by the horizontal dashed line) in IL-17A+ compared with IL-17A- CD8+ T-cells. Data are plotted as mean + SD and significance was analysed using paired Student’s t-test with Holm-Šídák multiple comparisons, adjusted p-values are reported. **(E, F)** After 3-day stimulation in the presence of hrIL-1β and hrIL-23, CD8+ T-cell cultures were re-stimulated for 3 hours with PMA/ionomycin and GolgiStop for identification of *in vitro*-induced IL-17A+ and IL-17A- CD8+ T-cell subsets by intracellular flow cytometry staining. Representative dot plots and cumulative data plotted as median + IQR show surface expression of CCR6 **(E)** and CD161 **(F)** on IL-17A+ or IL-17A- cells gated within CD8+ T-cells (n=8-10). Statistical analysis performed using Wilcoxon matched-pairs signed rank test.

We identified 25/29 IL-17A+ CD8+ T-cell signature genes to be differentially expressed with a relative fold change >1.5 compared with IL-17A- CD8+ T-cells (**Figure 3D**) and observed *IL17A* and *IL17F* were the most significantly differentially regulated genes (9.18-fold and 5.86- fold, respectively, both p<0.0001). Expression levels of IL-23 and IL-1β cognate receptor subunits *IL23R* and *IL1R1* were also significantly increased (3.43-fold, p<0.0001; and 1.94- fold, p<0.05, respectively) in IL-17A+ compared with IL-17A- CD8+ T-cells. *ZBTB16* (encoding PLZF), which was identified as the leading PC1 parameter in our primary PCA analysis, was further revealed as a signature type-17 gene differentially expressed in *in vitro*-generated IL-17A+ CD8+ T-cells (3.29-fold, p<0.01).

We confirmed the molecular data for the type-17 T-cell markers CCR6 and CD161 at the protein level by flow cytometry, which showed that a high proportion of *in vitro*-induced IL- 17A+ CD8+ T-cells co-expressed CCR6 and CD161 on their surface (median 88.1% and 93.4%, respectively), as compared with a much lower proportion of IL-17A- CD8+ T-cells (median 10.6% and 18.7%, respectively) (**Figure 3E, F**).

### Both IL-17-expressing CD8+ conventional T-cells and unconventional MAIT cells are induced *in vitro* upon type-17 polarisation

The transcriptional profiling revealed that *in vitro*-generated IL-17A+ CD8+ T-cells are endowed with prototypical type-17 markers however, several genes including *ZBTB16, ABCB1* (encoding multidrug resistance 1 (MDR-1))*, IL12RB2, IL18R1, DPP4* (encoding CD26)*, GZMB* and *KLRB1*, together with constitutively high surface expression of CD161, are associated with MAIT cell identity. This prompted us to investigate whether a proportion of *in vitro*-generated IL-17A+ CD8+ T-cells were comprised of MAIT cells. We stimulated PBMC with anti-CD3/CD28 mAbs in the absence or presence of IL-1β and IL-23 and assessed the expression of the MAIT cell-associated TCRVα7.2. Frequencies of IL-17A+ and IL-17F+ cells within Vα7.2- and Vα7.2+ CD8+ T-cells were quantified by intracellular cytokine staining. This analysis showed that IL-17A+ and IL-17F+ T-cells were significantly increased under type-17 polarising conditions within both Vα7.2- and Vα7.2+ CD8+ T-cells as compared with control conditions (**Figure 4A-C**). Notably, in all twelve donors tested, the frequencies of IL-17A and IL-17F expressing cells were significantly greater within Vα7.2+ compared with Vα7.2- CD8+ T-cells (median IL-17A+ 0.56% versus 0.18%; IL-17F+ 0.46% versus 0.16%, both p=0.002).

**Figure 4.**
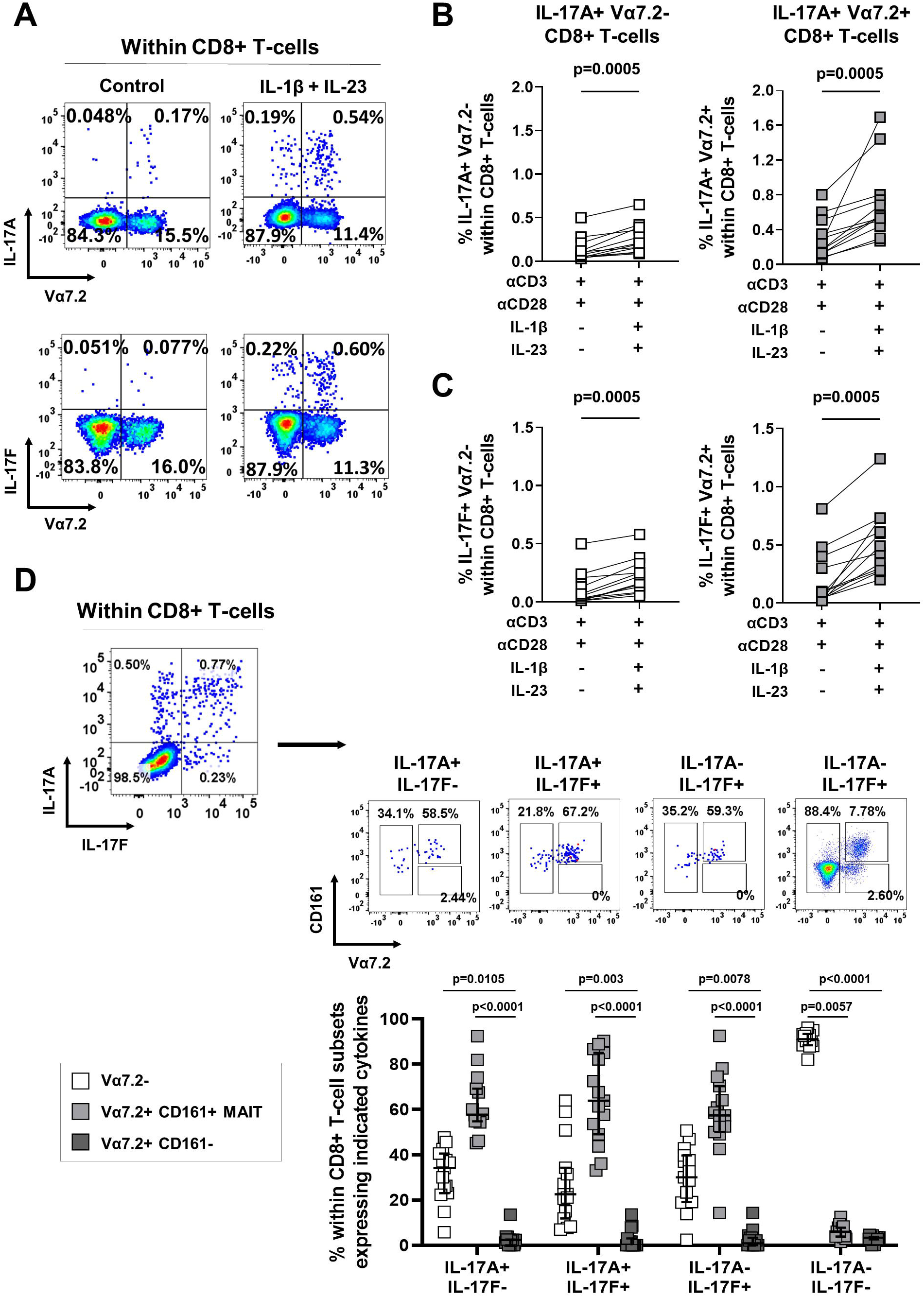
Type-17 *in vitro* polarisation increases the frequencies of both conventional and unconventional Vα7.2+ IL-17A+ CD8+ T cells. Healthy donor PBMC were cultured with soluble anti-CD3/CD28 mAbs in the absence or presence of hrIL-1β and hrIL-23 for 3 days, then re-stimulated for 3 hours with PMA/ionomycin and GolgiStop for assessment of intracellular IL-17A and IL-17F cytokine expression by flow cytometry. Representative staining plots **(A)** and cumulative data **(B, C)** show frequencies of IL-17A+ **(A, B)** or IL-17F+ **(A, C)** cells within conventional Vα7.2- (white squares) and unconventional Vα7.2+ (grey squares) CD8+ T-cells (n=10). **(D)** Representative dot plots showing the identification of IL-17A+IL- 17F-, IL-17A-IL-17F+, IL-17A+IL-17F+ and IL-17A-IL17F- cells within total CD8+ T-cells after culture in the presence of anti-CD3/CD28 stimulation with hrIL-1β and hrIL-23, and the proportions of CD161 and/or Vα7.2 expressing cells in each of these subsets. Cumulative data (n=15) are plotted as median + IQR. Statistical analysis was performed using Friedman test with Dunn’s multiple comparison.

Recent literature has shown that TCRVα7.2 bearing T-cells that do not express CD161 are transcriptionally distinct from CD161^hi^ Vα7.2+ MAIT cells (41). We therefore sought to further characterise the Vα7.2+ CD8+ T-cell compartment induced under type-17 polarising conditions by inclusion of CD161 co-stain. Within IL-17A and/or IL-17F expressing CD8+ T-cells, three populations were defined by TCRVα7.2 and CD161 expression: conventional Vα7.2- T-cells, Vα7.2+ CD161+ MAIT cells and Vα7.2+ CD161- T-cells. Frequencies of MAIT cells were the most abundant in *in vitro*-induced IL-17A+IL-17F-, IL-17F+IL-17A- and IL- 17A+IL-17F+ cells (median 57.7%, 58.6% and 63.5%) followed by conventional Vα7.2- CD8+ T-cells (median 34.1%, 28.6%, and 22.3%) (**Figure 4D**). In contrast, IL-17A and/or IL-17F producing cells only rarely contained Vα7.2+ CD161- CD8+ T-cells (median 2.4%, 0.9%, and 0%). We confirmed this finding using the MR1-5-OP-RU tetramer, which unequivocally identifies MAIT cells (42). Again, we found that upon type-17 polarisation IL-17A+ and IL- 17F+ cells were induced in both MR1-tetramer^neg^ conventional CD8+ T-cells and in MR1- tetramer^pos^ MAIT cells, with a statistically significant dominance of MAIT cells amongst the IL- 17-expressing cells (**Supplementary Figure 5**). We investigated if the preferential expansion of IL-17A+ cells within the MAIT cells was due to increased proliferation of MAIT cells during the culture period using CTV and Ki67 staining, however no substantive differences were observed between the MAIT and conventional CD8+ T-cell populations (**Supplementary Figure 6**).

### *In vitro*-induced IL-17A+ Va7.2- and IL-17A+ Va7.2+ CD8+ T-cells share a polyfunctional, pro-inflammatory phenotype

Having identified that unconventional Vα7.2+ MAIT cells represented around 60% of IL-17A and/or IL-17F expressing CD8+ T-cells within *in vitro*-induced cultures, we sought to compare whether the IL-17A+ Vα7.2- conventional and Vα7.2- MAIT CD8+ T-cell subsets displayed similar cytokine expression profiles. Comparable frequencies of IL-17A+ Vα7.2- and IL-17A+ Vα7.2+ CD8+ T-cells co-expressed GM-CSF (median 22.8% versus 24.5%), IL-17F (median 48.6% versus 43.6%) and IFNy (median 73.8% versus 78.6%), with only the frequency of TNFα significantly higher in the IL-17A+ Vα7.2+ compared with IL-17A+ Vα7.2- CD8+ T-cell subset (median 80.1% versus 89.9%, p=0.0313) (**Figure 5A**). Applying a Boolean gating strategy and the visualisation software SPICE, we explored the mono- and polyfunctional cytokine profiles of IL-17A+ Vα7.2- and IL-17A+ Vα7.2+ CD8+ T-cells (**Figure 5B, C**). This revealed that both *in vitro*-induced IL-17A+ CD8+ T-cell subsets predominantly comprised polyfunctional cells that express multiple pro-inflammatory cytokines (triple-positive for IL- 17A+IFNγ+TNFα+, orange pie segment, or quadruple-positive for IL-17A+IFNγ+TNFα+GM-CSF+, red pie segment). We only observed a statistically significant difference in the proportion of single-positive IL-17A+ cells which was higher in IL-17A+ Vα7.2- compared with IL-17A+ Vα7.2+ CD8+ T-cells (purple pie segment, p=0.0104) (**Figure 5B, C**). Additionally, in line with our molecular profile, we found that a high proportion of *in vitro*-generated IL-17A+ Vα7.2- as well as Vα7.2+ CD8+ T-cells harboured intracellular protein stores of granzyme B (n=2, median 94.6% versus 97.9%, respectively) and to a lesser extent granzyme A (n=2, median 49.1% and 69.4%, respectively) suggestive that both cell types also shared cytotoxic potential (data not shown).

**Figure 5.**
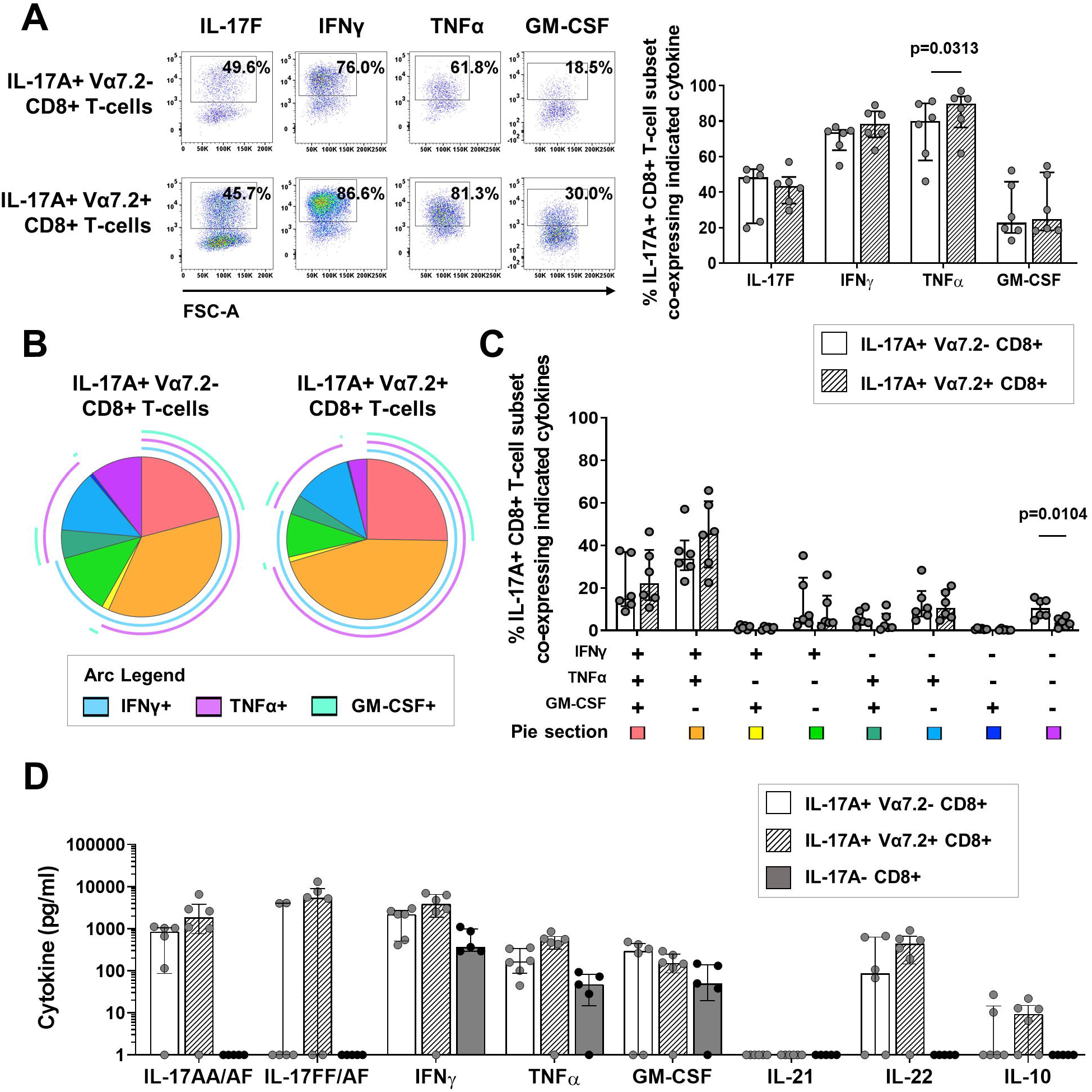
*In vitro*-induced IL-17A+ Vα7.2- and IL-17A+ Vα7.2+ CD8+ T-cell subsets have a similar type-17 cytokine profile. Bulk CD8+ T-cells isolated from healthy donor PBMC were cultured for 3 days with plate-bound anti-CD3 mAb and soluble anti-CD28 mAb in the presence of hrIL-1β and hrIL-23. On day 3, cells were re-stimulated for 3 hours with PMA/ionomycin and GolgiStop for assessment of intracellular cytokine expression. **(A)** Representative flow cytometric staining and cumulative data (n=6) show the frequencies of IL-17A+ Vα7.2- (white bars) and IL-17A+ Vα7.2+ (hashed bars) CD8+ T-cell subsets that express IL-17F, IFNγ, TNFα or GM-CSF. **(B, C)** Polyfunctional cytokine data were analysed using SPICE software. Pie charts **(B)** depict cytokine profiles of IL-17A+ Vα7.2- (left) and IL- 17A+ Vα7.2+ (right) CD8+ T-cells with the individual and overlapping arcs indicating the double, triple or quadruple cytokine combinations produced by each proportion of cells. **(C)** Cumulative frequencies of each pie section within IL-17A+ Vα7.2- (white bars) and IL-17A+ Vα7.2+ (hashed bars) are plotted as median + IQR (n=6). **(D)** CD8+ T-cells were FACS sorted using an IL-17A cytokine secretion assay into IL-17A+ Vα7.2- (white bars), IL-17A+ Vα7.2+ (hashed bars) or IL-17A- (grey bars). Cells were cultured for 20 hours and supernatants collected for detection of IL-17AA/AF, IL-17FF/AF, IFNγ, TNFα, GM-CSF, IL-21, IL-22 and IL- 10 by Luminex. Statistical analysis performed using Wilcoxon matched-pairs signed rank test.

Using a Luminex^®^ assay, we then measured the production of the cytokines IL-17AA/AF, IL- 17FF/AF, IFNγ, TNFα, GM-CSF, IL-21, IL-22 and IL-10 by sorted IL-17A+ Vα7.2-, IL-17A+ Vα7.2+or total IL-17A- CD8+ T-cell subsets (representative gating strategies shown in **Supplementary Figures 4A and D**). Both conventional Vα7.2- and unconventional Vα7.2+ IL-17A+ CD8+ T-cells showed production of IL-17AA/AF, IFNγ and TNFα (**Figure 5D**). More variation was observed in the production of IL-17FF/AF, GM-CSF, IL-22 and IL-10, with Vα7.2+ IL-17A+CD8+ T-cells showing more consistent production of these cytokines than Vα7.2- IL-17A+CD8+ T-cells. Neither subset produced IL-21. *In vitro-induced* IL-17A- CD8+ T-cell cultures were assessed as a negative control, which showed no production of the type- 17 associated cytokines IL-17AA/AF, IL-17FF/AF, IL-21 and IL-22 but rather secreted IFNγ, TNFα and GM-CSF only.

### *In vitro*-induced IL-17A+ CD8+ T-cells are functional, with capacity to induce pro-inflammatory cytokine production by PsA synovial fibroblasts

As our final step, we sought to determine the functional contribution of *in vitro*-induced IL- 17A+ CD8+ T-cells, by investigating their ability to promote clinically relevant pro-inflammatory cytokine production in an *in vitro* model of joint inflammation. For this, cell culture supernatants were collected from *in vitro*-induced and then CSA-FACS-sorted IL-17A+ or IL- 17A- CD8+ T-cells. Supernatants were added (20% v/v) to synovial tissue fibroblasts from patients with PsA. After 24 hours, fibroblast culture supernatants were collected for analysis of the pro-inflammatory cytokines IL-6 and IL-8. Addition of IL-17A+ CD8+ T-cell culture supernatant led to a significant increase in IL-6 and IL-8 production by PsA fibroblasts as compared with fibroblasts cultured in medium alone, whilst no significant increase in either IL-6 or IL-8 secretion was observed when PsA fibroblasts were cultured in the presence of IL-17- CD8+ T-cell supernatants (**Figure 6A**).

**Figure 6.**
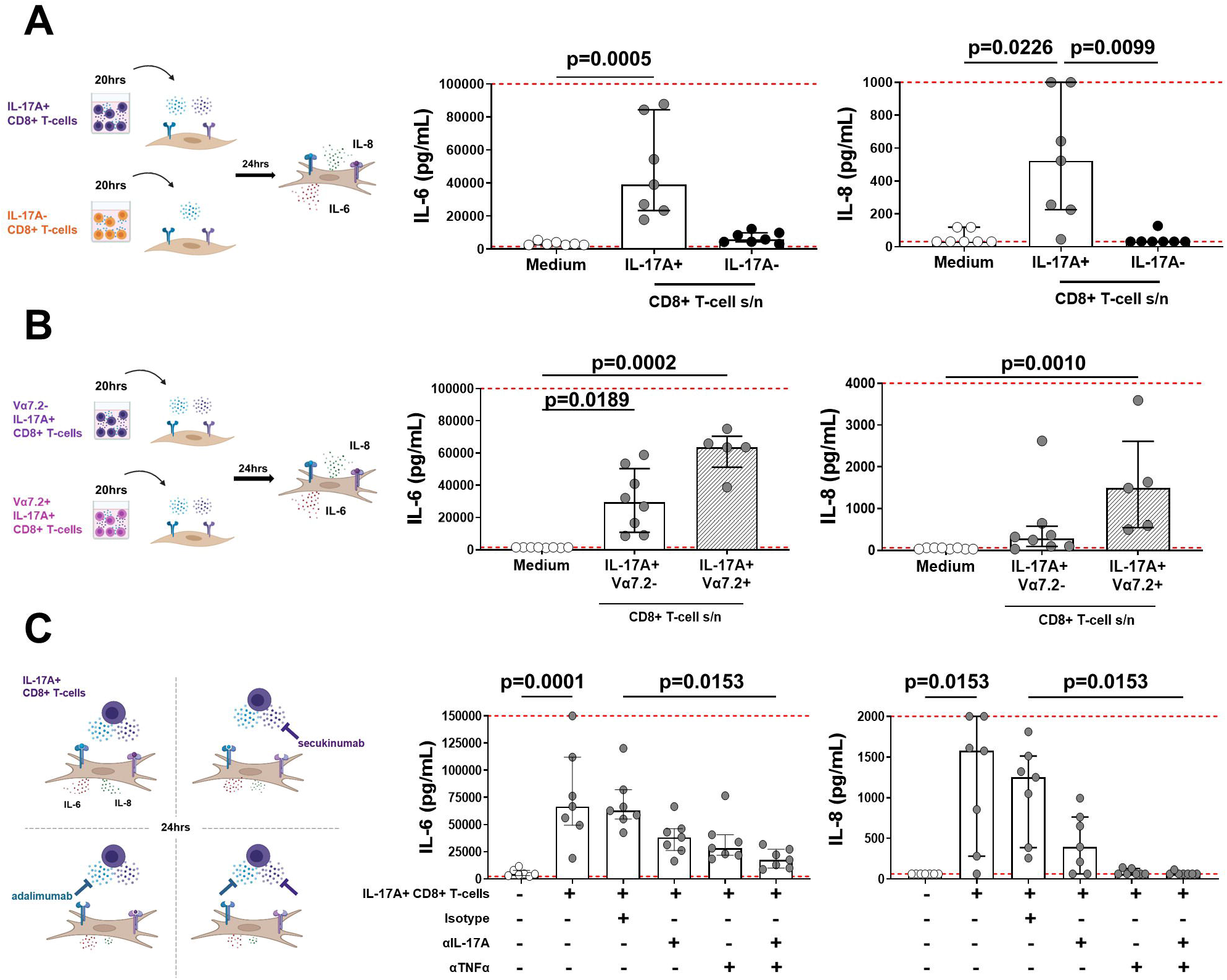
*In vitro*-induced IL-17A+ CD8+ T cells are biologically functional. Bulk IL-17A+ and IL-17A- CD8+T-cells or IL-17A+ Vα7.2- and IL-17A+ Vα7.2+ CD8+ T-cells were FACS sorted using an IL-17A CSA following 3-day culture with plate-bound anti-CD3, soluble anti- CD28 mAbs in the presence of hrIL-1β and hrIL-23. The T-cells or their post-sort 20-hour cell culture supernatants (20% v/v) were added to PsA synovial fibroblasts (1×10^4^) for 24 hours. IL-6 (left panels) and IL-8 (right panels) production in fibroblast culture supernatants were measured by ELISA. Schematics depict experimental workflows. Cumulative data show IL-6 and IL-8 production from fibroblasts cultured with **(A)** supernatants from sorted IL-17A+ or IL- 17A- CD8+ T-cells, **(B)** supernatants from IL-17A+ Vα7.2- or IL-17A+ Vα7.2+ CD8+ T-cells or **(C)** total IL-17A+ CD8+ T-cells (2.5×10^4^) in the absence or presence of isotype control, anti-IL-17A or anti-TNFα neutralising antibodies. Data are plotted as median + IQR with dashed red lines indicating lower and upper ELISA detection limits. Statistical analysis was performed using a matched-paired Friedman test **(A, C** n=7) or unmatched Kruskal Wallis test **(B** n=5-7) with comparison to medium **(A-C)** and isotype **(C)** by Dunn’s Multiple Comparisons test.

We next aimed to determine whether there were differences in these pro-inflammatory responses when fibroblasts were cultured in the presence of secretory products from *in vitro*-induced IL-17A+ Vα7.2- conventional or IL-17A+Vα7.2+ MAIT CD8+ T-cells. Addition of supernatants from either cell type induced a significant increase in IL-6 production compared with fibroblasts cultured in medium alone. IL-8 levels were significantly increased in the presence of cell culture supernatant from IL-17A+ Vα7.2+ CD8+ T-cells but not from IL-17A+ Vα7.2- CD8+ T-cells. Whilst the median IL-6 and IL-8 production was higher in presence of cell culture supernatant from IL-17A+ CD8+ MAIT cells compared with conventional T-cells (median 63,700pg/mL versus 29,500pg/mL and 1,500pg/mL versus 280pg/mL, respectively), this difference did not reach statistical significance (**Figure 6B**).

IL-17A is known to act synergistically with TNFα to promote pro-inflammatory cytokine production by stromal cells (4,5). Given our observation that *in vitro*-induced IL-17A+ CD8+ T-cell subsets actively secreted IL-17A and TNFα (**Figure 5**), we investigated the effect of single and dual blockade of these cytokines in co-cultures of PsA fibroblasts and FACS-sorted *in vitro*-induced IL-17A+ CD8+T-cells (1:2.5 cell ratio). First, we identified that as with T-cell supernatants, fibroblasts co-cultured with *in vitro*-induced IL-17A+ CD8+ T-cells produced significantly higher levels of IL-6 and IL-8 compared with fibroblast monocultures (p=0.0001 and p=0.0153, respectively) (**Figure 6C**). Addition of an isotype control mAb to the co-cultures did not affect IL-6 or IL-8 production. IL-6 and IL-8 production was lower in the presence of either secukinumab (anti-IL-17A) or adalimumab (anti-TNFα) and was significantly reduced upon combined blockade of IL-17A and TNFα (p=0.0153). Collectively, these data demonstrate that *in vitro*-induced IL-17A+ CD8+ T-cells and their secretory products are biologically active, with capacity to significantly increase pro-inflammatory IL-6 and IL-8 production by synovial fibroblasts from patients with PsA.

## Discussion

We report herein that human IL-17A+ and/or IL-17F+ CD8+ T-cells can be expanded *in vitro* upon anti-CD3 and anti-CD28 stimulation in the presence of IL-1β and IL-23. These cells are characterised by typical type-17 phenotype, cytokine and transcriptional profiles. Furthermore, we demonstrate that *in vitro* polarisation induces both conventional IL- 17A+CD8+ T-cells as well as unconventional IL-17A+ CD8+ MAIT cells, but that these cell types have a shared phenotype and are functionally active, with the capacity to drive biologically relevant pro-inflammatory cytokine production from PsA synovial tissue-derived stromal cells.

Our study confirms previous reports that detected low *ex vivo* frequencies of IL-17A+ CD8+ T-cells in healthy human peripheral blood with a rare presence of IL-17F+ CD8+ T-cells (40, 43–46) and adds weight to the few studies that reported on the expansion of human IL-17- expressing CD8+ T-cells (22, 23). We also demonstrate that equivalent IL-17A+ CD8+ T cell induction can be achieved using either cryopreserved or freshly isolated PBMC, indicating that our protocol can be used with bio-banked samples. Kondo et al previously showed a limited percentage of Tc17 cells (0.11%) were differentiated upon culture of human naïve CD8+ T-cells with anti-CD3/CD28 mAbs in the presence of TGFβ, IL-6, IL-1β and IL-23 for 5 days and supplemented with IL-2 for a further 4 days (22). Gras et al instead stimulated bulk CD8+ T-cells with anti-CD3/CD28 mAbs, TGFβ and IL-6 for 3 days and measured secreted IL-17A and IL-17F in culture supernatants by ELISA as readouts of Tc17 induction (23). The authors reported elevated secretion of IL-17A and IL-17F compared with the control condition (anti-CD3/CD28 stimulation without TGF-b and IL-6). These studies each tested a very small sample size (n=2-3) and presented limited assessment on either the frequency or immunophenotype of IL-17A and/or IL-17F+ CD8+ T-cells. Our *in vitro* protocol exceeds those previously detailed since we demonstrate that culturing either PBMC or purified CD8+ T-cells for 3-days with anti-CD3/CD28 stimulation in the presence of IL-1β and IL-23 yields a significant, and on average 3-5-fold increase in the frequencies of IL-17A+ CD8+ T-cells compared with anti-CD3/CD28 stimulated cultures alone. Additionally, we demonstrate a significant, 7-fold increase in IL-17F+ CD8+ T-cells when PBMC were stimulated in the presence of IL-1β and IL-23. Recent studies have detailed the use of *in vitro* polarising cultures to investigate pharmacological or cytokine directed modulation of Tc17 cell responses (13, 26) yet, as with earlier literature, their polarising protocols are not directly comparable. Li et al. reported a 20-fold increase in Tc17 cell frequency of healthy and colorectal cancer patient naïve CD8+ T-cell cultures upon 7-day culture in IMDM media using anti-CD3/CD28 mAb stimulation with IL-2, IL-1β, IL-6, IL-23, TGFβ, anti-IL-4 and anti-IFNγ (26). Whilst Globig et al. reported a maximum induction of ~0.6% IL-17A+ CD8+ T-cells within healthy donor PBMC cultured with T-cell activator in the presence of IL-1β, IL-6, IL-23 and TGFβ (13). Neither study performed extensive Tc17 cell immunophenotyping post-culture as we report here. Compared with the aforementioned human Tc17 protocols, our data from purified CD8+ T-cell cultures suggest IL-1β and IL-23 were sufficient to induce an increase in IL-17 expression in human CD8+ T-cells. When our polarising condition was supplemented with IL-6 or anti-IFNγ, we did not observe consistently increased frequencies of IL-17A+ CD8+ T-cells. This is in line with previous studies that showed that IL-6 is not a requirement in human Th17 cell differentiation and that IFNγ neutralisation could either promote or antagonise Th17 cell polarisation depending on the timing of administration (47, 48).

Transcriptional analysis revealed that our *in vitro* polarising culture confers a strong type-17 signature in induced IL-17A+ CD8+ T-cells distinct from IL-17A- CD8+ T-cells. This signature was characterised by high expression of type-17 related genes including *IL17A, IL17F, RORC, RORA, MAF, CCR6, CXCR6, KLRB1 and IL23R*, which is concordant with previous immunophenotyping reports of human Tc17 cells in health and at sites of inflammation (18, 40, 49). Additionally, some of the more novel markers associated with human Th17 and/or Tc17 cells were found more abundantly expressed in our IL-17A+ CD8+ T-cells including *CTSL, HOPX, MCAM* and *GPR65* (encoding cathepsin L, homeodomain-only protein homeobox, melanoma cell adhesion molecule and G protein-coupled receptor 65, respectively) (50–54); as was *CTLA4* (encoding cytotoxic T-lymphocyte associated protein 4) which has been described as a regulator of murine Tc17 cell differentiation and stability (55). We were unable to conclude on the expression of *IL17B, IL17C, IL17D* or *IL17E* transcript in IL-17A+ CD8+ T-cells as these were filtered from downstream statistical assessments due to normalised expression lower than that of *CD4*. We also observed a degree of molecular similarity in both sorted subsets with similar mRNA expression of surface receptors (*IL6R, IL18R1, IL21R*), transcription factors (*TBX21, EOMES, STAT3, IRF4, BCL11B*) and effector molecules (*IFNG, TNF*). We confirmed by flow cytometry that *in vitro*-induced IL-17A+ CD8+ T-cells predominantly co-expressed the hallmark type-17 surface markers CCR6 and CD161.

A small proportion of IL-17A- CD8+ T-cells also expressed CCR6 and CD161 (also at lower transcript levels than IL-17A+ counterparts), which is consistent with literature highlighting that these surface markers are not exclusively restricted to human IL-17 expressing T-cells (38, 56). This finding reiterates the need for identification of additional lineage specific surface markers to better facilitate identification of Tc17 cells and type-17 cells more broadly. Taken together, these analyses validate the cytokine secretion assay approach for specifically purifying type-17 cells that secrete bioactive IL-17A.

The leading transcriptional parameter defining segregation of IL-17A+ from IL-17A- CD8+ T-cells was identified as PLZF, which is commonly associated with directing type-17 effector function in unconventional rather than conventional T-cells. With further immunoprofiling, our analysis indeed revealed that *in vitro* polarisation did not only induce IL-17A and IL-17F expression in conventional CD8+ T cells, but also in Vα7.2+/MR1-tetramer^pos^ MAIT cells. This finding is in line with reports evidencing that IL-17A and IL-17F production by MAIT cells requires cooperative TCR signalling with specific cytokine signals, such as IL-1β and IL-23 (37, 49), IL-7 (35), or IL-12 with IL-18 (31, 57). Interestingly, the largest fraction of IL-17A/F expressing cells was contained within the MAIT population. One explanation could be that there was preferential expansion of IL-17A/F-producing MAIT cells upon *in vitro* culture, however our preliminary data did not reveal substantive differences in proliferative capacity between IL-17A+ CD8+ MAIT and conventional T-cells.

It was interesting to note that our PBMC polarising conditions led to a relative greater fold increase in IL-17F than IL-17A (7-fold versus 3-fold). Additionally, both conventional and MAIT IL-17A+ CD8+ T cells comprised IL-17A or IL-17F single and double positive cells. We have previously demonstrated that CD28 costimulation differentially regulates IL-17F versus IL-17A expression in CD4+ T-cells (5). Furthermore, a previous study showed a 10-fold higher production of IL-17F compared with IL-17A upon anti-CD3/CD28 stimulation of CD8+ T-cells (23), and IL-17F was found to be the dominant isoform when MAIT cells were stimulated in the presence of IL-12 and IL-18 (57). Collectively, these findings suggest differential regulation of IL-17A and IL-17F production in human T-cells.

Even though *in vitro* polarisation induced two distinct populations of IL-17A+ CD8+ T cells, our data indicate that Vα7.2- conventional and Vα7.2+ MAIT IL-17A+ CD8+ T-cell types share a common polyfunctional profile and actively secrete a combination of pro-inflammatory cytokines including IL-17A, IL-22, GM-CSF, IFNγ and TNFα. Of note, IL-17A+ Vα7.2+ CD8+ T-cells appeared to induce a more potent pro-inflammatory response in PsA-derived synovial fibroblasts, on average inducing greater levels of IL-6 and IL-8 compared with IL-17A+ Vα7.2- conventional CD8+ T-cells. This may be attributable to elevated levels of TNFα, and in some donors more IL-17FF/AF, in IL-17A+ Vα7.2+ CD8+ T-cells compared with the IL-17A+ Vα7.2- CD8+ T-cell counterparts. These subtle differences in our findings strengthen the importance to distinguish between classical and unconventional cell types, which to date few studies have performed when broadly identifying IL-17-producing CD8+ T-cells. This may indeed explain research discrepancies and for future assessments, it is essential protective or pathological contributions of individual subsets are clearly ascertained.

By addition of secukinumab and adalimumab in IL-17A+CD8+ T-cell–fibroblast co-cultures, we reaffirmed cooperation of IL-17A with TNFα in driving pro-inflammatory cytokine production in the context of PsA. Whilst IL-17F has reduced potency relative to IL-17A, we previously showed IL-17F can similarly synergise with TNFα to elicit significant inflammatory responses in synovial fibroblasts from PsA and RA patients (5). Dual blockade of IL-17A and IL-17F by bimekizumab also more effectively reduced IL-17-driven secretion of IL-6 and IL-8 by synovial fibroblasts compared with blockade of IL-17A alone (5, 57, 58). An IL-17F CSA was not commercially available, however it would be of interest to isolate IL-17A and IL-17F single and double positive T-cell subsets to investigate further their individual versus combined functional roles.

We verified functionality of our *in vitro*-generated IL-17A+ CD8+ T-cells and for the first-time report clinically relevant modulation of IL-17A+ CD8+ T-cell-PsA synovial fibroblast interactions, which suggests a pro-inflammatory contribution of Tc17 cells to PsA joint inflammation. A pathogenic function of human Tc17 cells has previously been demonstrated in adoptive transfer models. Hu et al. showed that adoptive transfer of human *in vitro*-generated CAR-directed type-17 T-cells (a mixture of Th17 and Tc17) primed with a RORγt agonist mediated potent anti-tumour responses in a murine melanoma model (24). Mechanistically, they showed that co-transfer of Th17 cells with Tc17 cells mediated robust and long-lived anti-tumour immunity, consistent with previous publications which showed that Th17 cells can augment the activation of CD8+ T-cells (59, 60). Hinrichs et al. also reported that type 17-polarised CD8+ T-cells mediated enhanced anti-tumour immunity and demonstrated greater persistence than non-polarised CD8+ T-cells (61). An in-depth study by Gartlan et al., using fate mapping reporter mice also showed that mouse Tc17 cells differentiate during GVHD culminating in a highly plastic, hyperinflammatory, poorly cytolytic effector population, which they termed “inflammatory iTc17” (62). Targeted depletion of these inflammatory iTc17 cells resulted in protection from lethal GVHD.

These *in vivo* data together with our *in vitro* data strongly suggests that Tc17 cells are biologically relevant contributors to inflammation in diseases where an enrichment of these cells is found. In addition, our *in vitro* induction approach has important potential to improve mechanistic understanding of how Tc17 cells contribute to pathogenic, as well as homeostatic, immune responses, which could offer novel translational insights into therapeutic targeting of these cells.

## Data Availability Statement

The original contributions presented in the study are included in the article/Supplementary Material. Further inquiries can be directed to the corresponding author.

## Author Contributions

LT, BK and KS contributed to study conception and supervision. LT, US and EG designed experiments that were equally performed, the data acquired, analysed and interpreted by US and EG. LD and SL aided with CSA-FACS sorting experiments and AC kindly provided some synovial tissue fibroblast lines for functional assessments. BK coordinated PsA patient recruitment and sample collection during clinic at the Rheumatology Department, Guy’s hospital. US wrote an initial draft of the manuscript, which was re-worked and finalized by EG with editing by LT. All authors except AC, who sadly passed away during completion of the project, were involved in critically revising the article and approved the final submitted version.

## Acknowledgments

The authors would like to express their thanks to the healthy volunteers and PsA patient participants, without whom the work would not have been possible. We are also grateful to Dr Rocio Martinez-Nunez, Dr Paul Lavender and Kate Poulton for their considered intellectual advice and input guiding molecular experimental design, and to Dasha Freydina for performing Bioanalyser assessments (all from King’s College London). The MR1 tetramer technology was developed jointly by J McCluskey, J Rossjohn and D Fairlie, and the material was produced by the NIH Tetramer Core Facility as permitted to be distributed by the University of Melbourne. Schematic figures were created with BioRender.com.

