## Supplementary Figures1-6. for "Phenotypic, molecular and functional characterisation of human *in vitro*-generated IL-17A+ CD8+ T-cells"

**Figure S1**

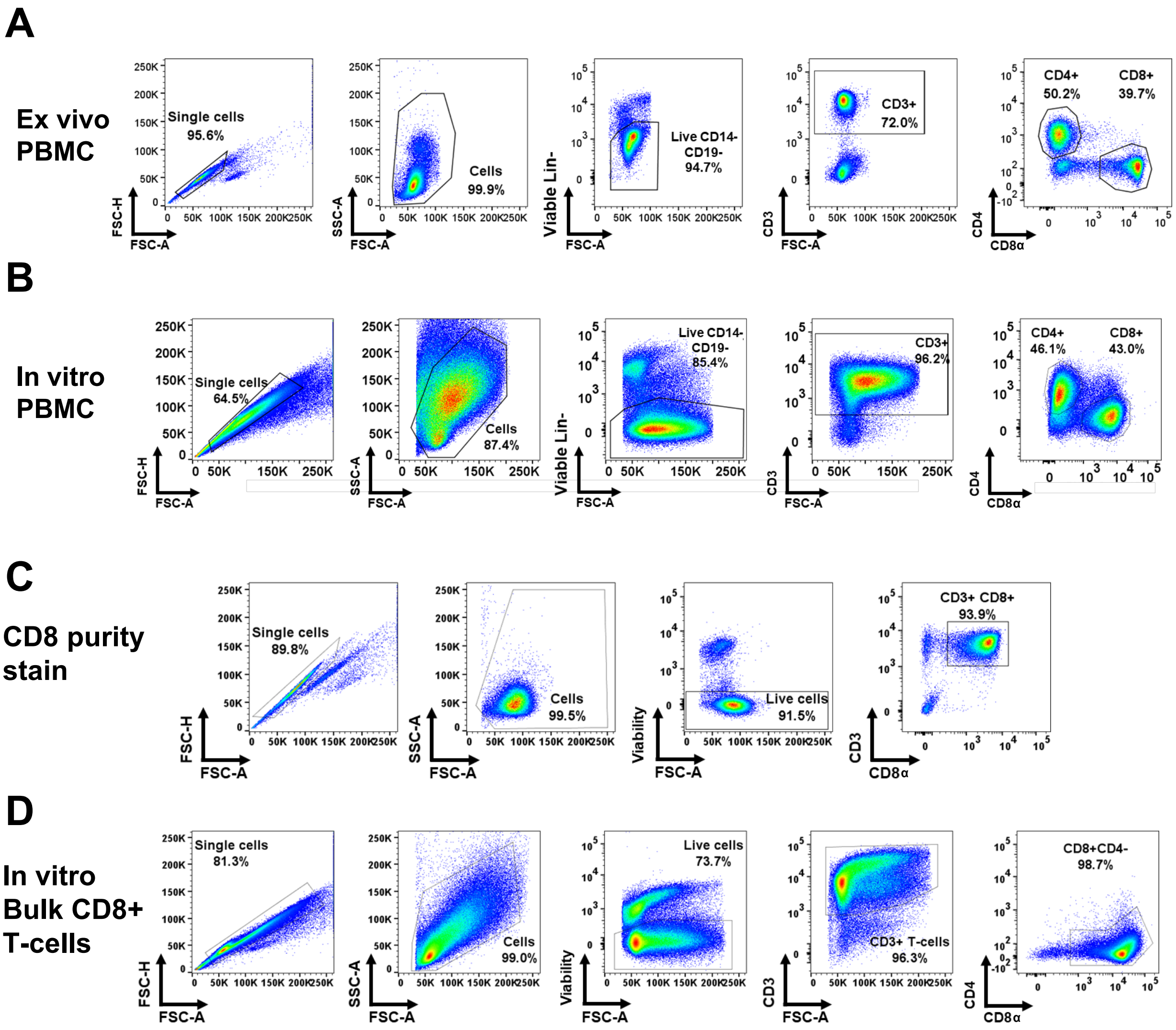


**Supplementary Figure 1. Gating strategies for CD8+ and CD4+ T-cell identification within PBMC ex vivo or following in vitro cell culture.** Healthy donor whole PBMC, either ex vivo **(A)** or following *in vitro* culture **(B)** were first gated on single cells and FSC/SSC. Dead cells and CD14+ cells were then excluded, followed by gating on CD3+ T-cells, from which CD8+ or CD4+ T-cell subsets were identified. **(C)** Representative purity staining for CD8+ T-cells isolated by magnetic bead separation from whole PBMC. Average CD8+ T-cell purity was 93% as determined by flow cytometry. **(D)** Gating strategy applied to identify live CD3+ CD8+ T-cells in CD8+ T-cell cultures following 3 day *in vitro* culture.

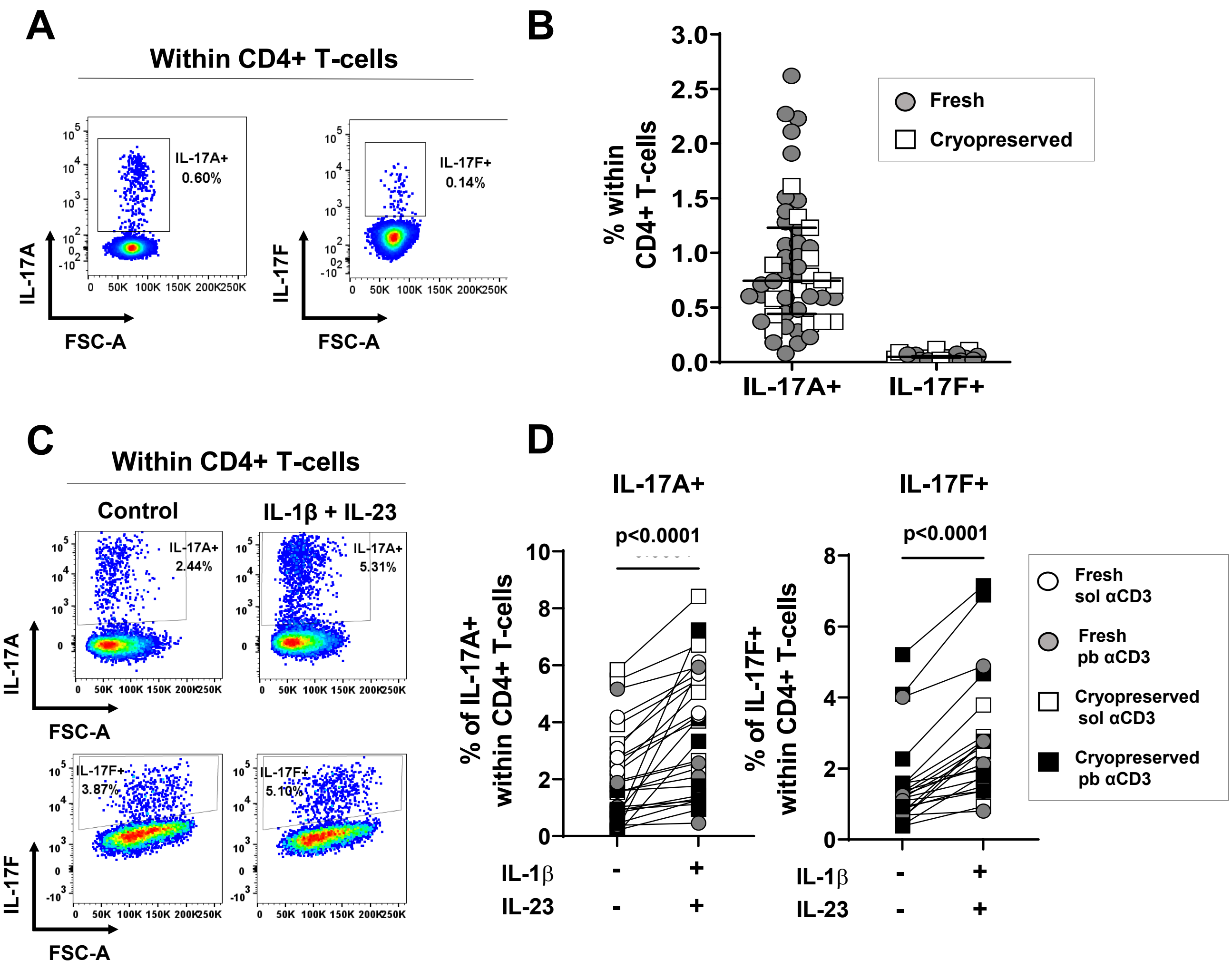


**Supplementary Figure 2. Type-17 polarising conditions promote IL-17A and IL-17F expressing CD4+ T-cells. (A, B)** Freshly isolated (circles) or cryopreserved (squares) healthy donor PBMC were stimulated ex vivo for 3 hours with PMA, ionomycin and GolgiStop for assessment of intracellular IL-17A and IL-17F cytokine expression by CD4+ T-cells using flow cytometry. Representative staining plots **(A)** and cumulative data **(B)** show frequencies of IL-17A+ and IL-17F+ cells within live CD3+ CD4+ T-cells from independent donors (n=50 and n=22, respectively). **(C, D)** Fresh (circles) or cryopreserved (squares) healthy donor PBMC were cultured for 3 days with either plate-bound (filled symbols) or soluble (open symbols) anti-CD3 mAb and soluble anti-CD28 mAb in the absence (control) or presence of hrIL-1β and hrIL-23. After 3 days cells were re-stimulated with PMA, ionomycin and GolgiStop for detection of intracellular cytokine expression by CD4+ T-cells. Representative staining plots **(C)** and cumulative data **(D)** show frequencies of IL-17A+ and IL-17F+ cells within live CD3+CD4+ T-cells from independent donors (n=27 and n=22, respectively). Statistical analysis performed using Wilcoxon matched-pairs signed rank test.

### Figure S3

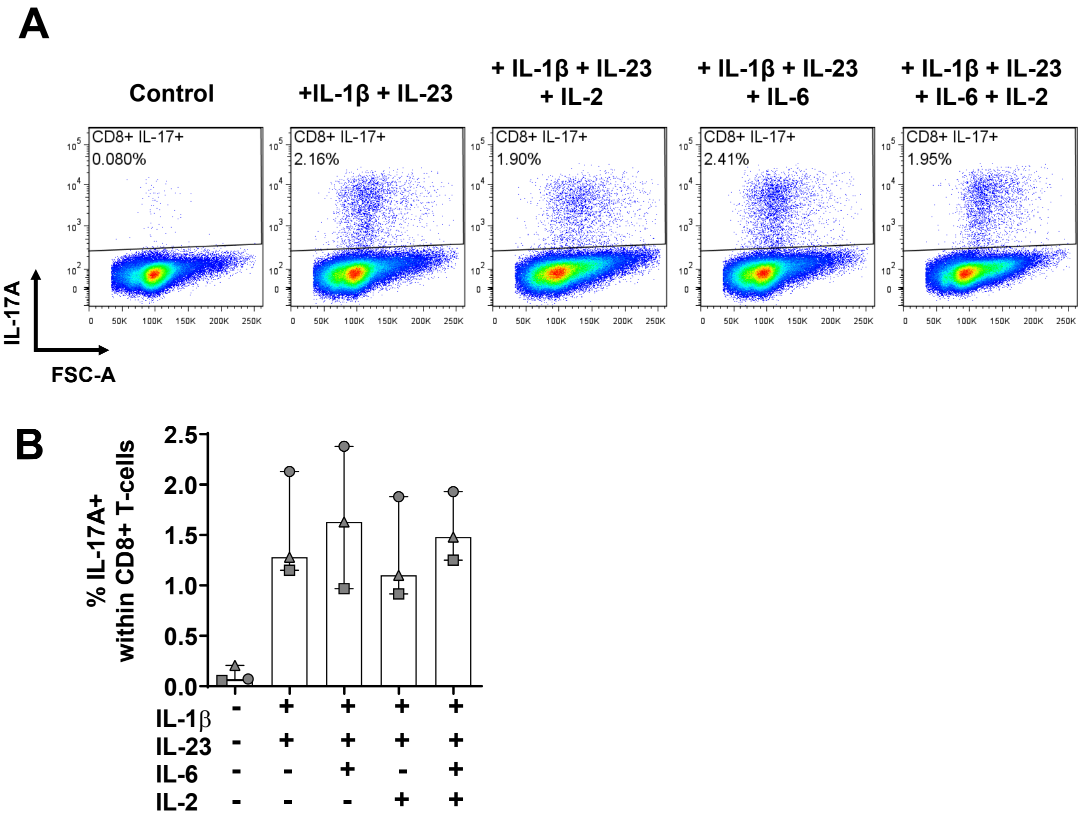


**Supplementary Figure 3. Addition of hrIL-6 and/or hrIL-2 to in vitro cultures in combination with IL-1β and IL-23 does not further increase the frequency of induced IL-17A+ CD8+ T-cells.** CD8+ T-cell cultures were cultured in the presence of anti-CD3/CD28 beads (1:10 bead to cell ratio) in the absence (control) or presence of hrIL-1β and IL-23 with or without rhIL-6 (20ng/mL) for 3 days. On day 3, media containing recombinant cytokines was replenished and supplemented with or without hrIL-2 and cells cultured for an additional 3 days. On day 6, cells were re-stimulated for 3 hours with PMA, ionomycin and GolgiStop and intracellular cytokine expression assessed by flow cytometry. **(A)** Representative staining plots and **(B)** cumulative data (n=3) showing the frequencies of IL-17A+ CD8+ T-cells across culture conditions. Data are plotted as median ± IQR.

### Figure S4

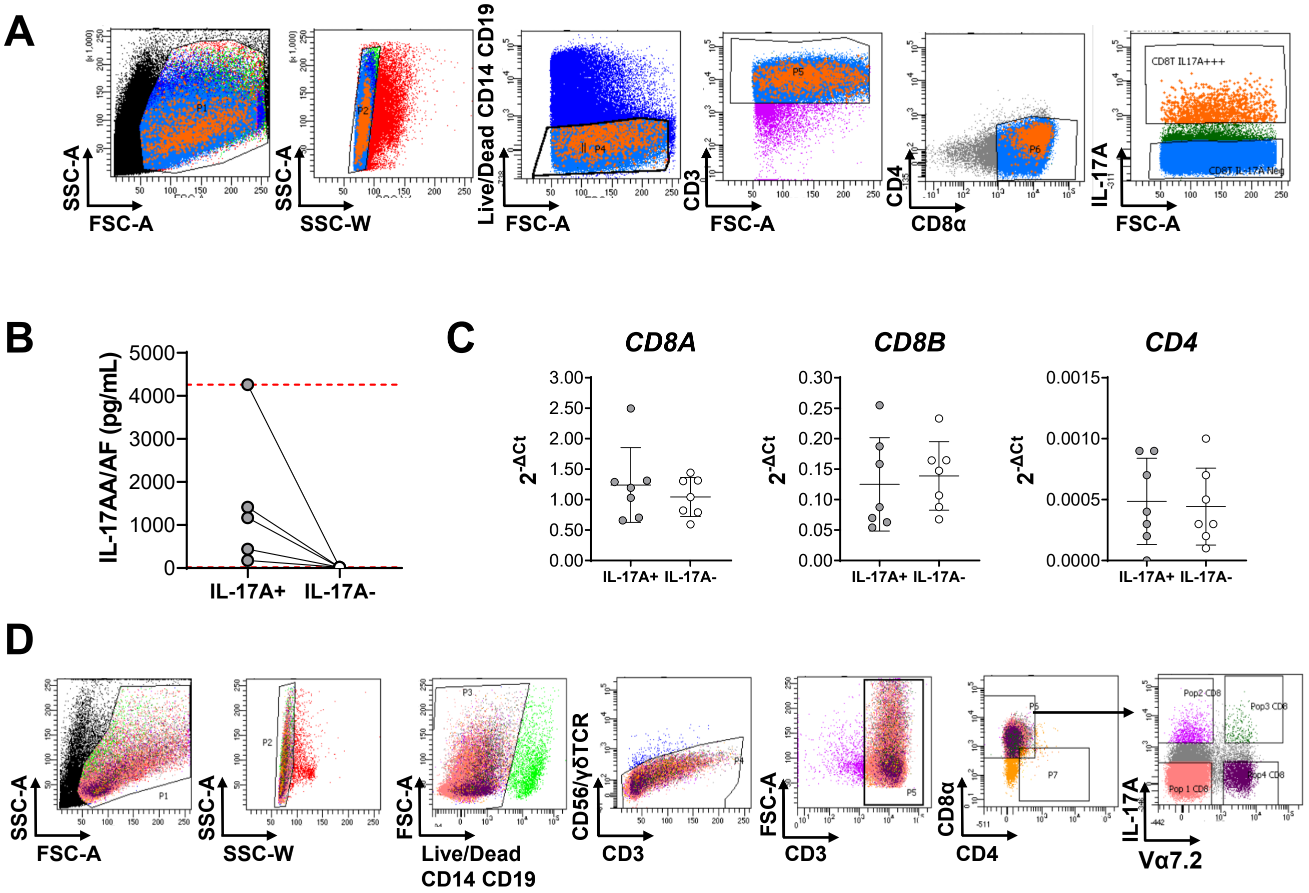


**Supplementary Figure 4. Sorting of IL-17A-secreting CD8+ T-cells using an IL-17A cytokine secretion assay.** Bulk CD8+ T-cells were cultured under type-17 polarising conditions for 3 days followed by 1.5 hours stimulation with PMA/ionomycin to allow for IL-17A secretion and capture using an IL-17A cytokine secretion assay. IL-17A secreting (IL-17A+) and non-secreting (IL-17A-) CD8+ T-cell subsets were then FACS sorted according to surface marker and IL-17A detection antibody staining. Representative gating strategies used to identify **(A)** IL-17A+ and IL-17A- CD8+ T-cells. **(B)** Sorted IL-17A+ and IL-17A- T-cells were cultured for 20 hours in culture medium to generate supernatants which were analysed for IL-17AA/AF by Luminex assay (n=5). **(C)** Dot plots show normalised mRNA expression levels of *CD8A, CD8B* and *CD4* within the indicated CD8+ T-cell populations as quantified by qPCR array (n=7 independent donors). Expression was normalised to geometric mean of endogenous genes *B2M* and *PPIA*. Gene expression is reported as mean ± SD. **(D)** Representative gating strategy used to identify IL-17A+ Vα7.2- and IL-17A+ Vα7.2+ populations by CSA-FACS sorting from bulk CD8+ T-cell type-17 polarised cultures re-stimulated on day 3 for 1.5hrs with PMA/ionomycin. Sorted IL-17A+ subsets were cultured for 20 hours in culture medium to generate supernatants for Luminex assay.

### Figure S5

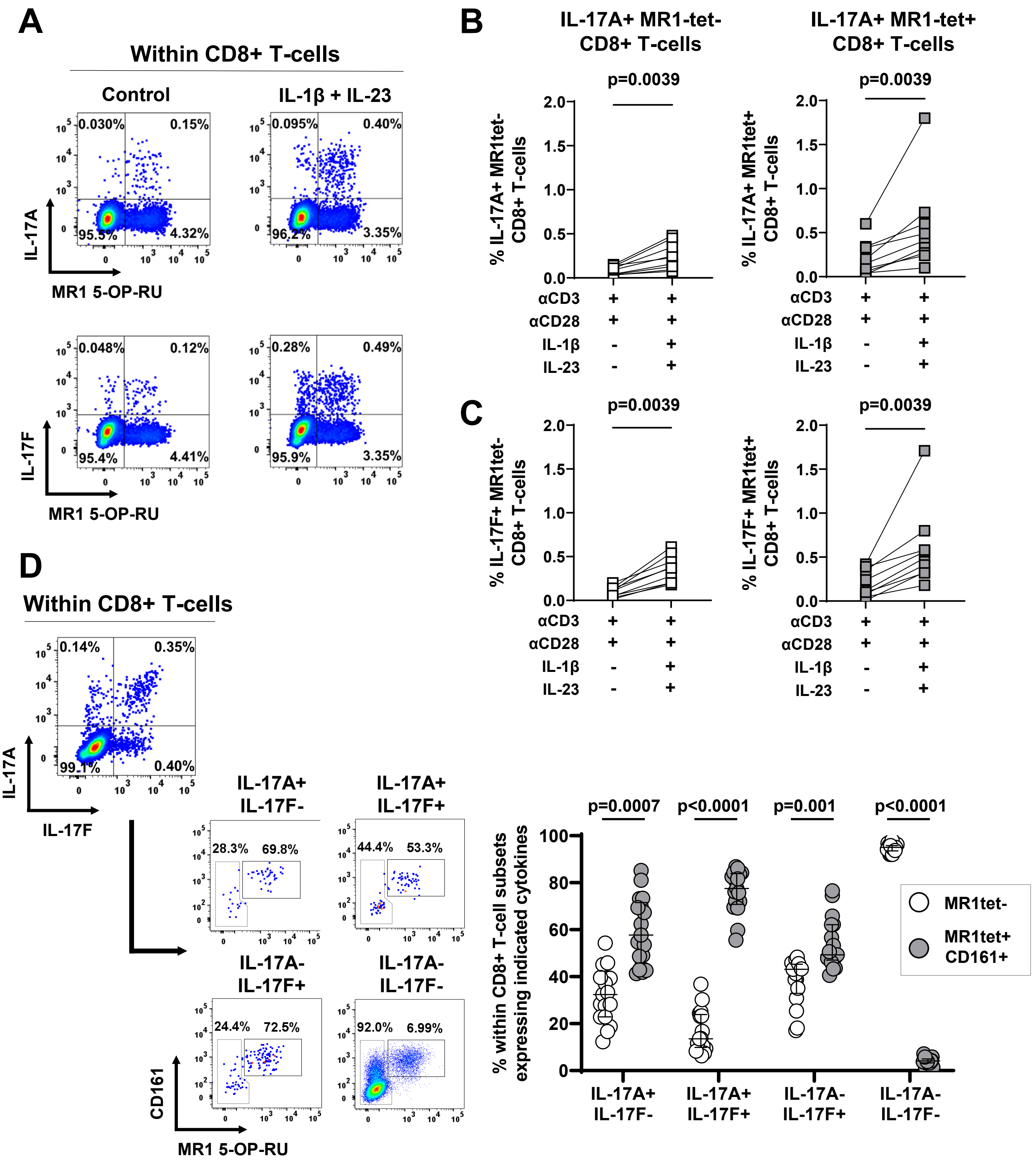


**Supplementary Figure 5. Both IL-17-expressing CD8+ conventional and unconventional MAIT cells are induced within type-17 polarising in vitro cultures.** Healthy donor PBMC were cultured for 3 days with plate-bound anti-CD3, soluble anti-CD28 mAbs in the absence or presence of IL-1β and IL-23 followed by 3 hours with PMA, ionomycin and GolgiStop. Representative staining plots **(A)** and cumulative data **(B, C)** show frequencies of IL-17A+ **(A, B)** or IL-17F+ **(A, C)** cells within MR1 5-OP-RU tetramer-negative (white squares) and tetramer-positive (grey squares) CD8+ T-cells (n=5). **(D)** Representative dot plots showing frequencies of IL-17A+IL-17F-, IL-17A+IL-17F+, IL-17A-IL-17F+ and IL-17A-IL17F- cells gated within total CD8+ T-cells after culture in the presence of anti-CD3/CD28 stimulation with IL-1β and IL-23, and the subsequent proportions of MAIT cells (identified by MR1 5-OP-RU tetramer and CD161 positive cells) that comprise each IL-17 cytokine expressing population. Cumulative data (n=17) are plotted as median ± IQR and statistical analysis performed using Wilcoxon matched-pairs signed rank test.

### Figure S6

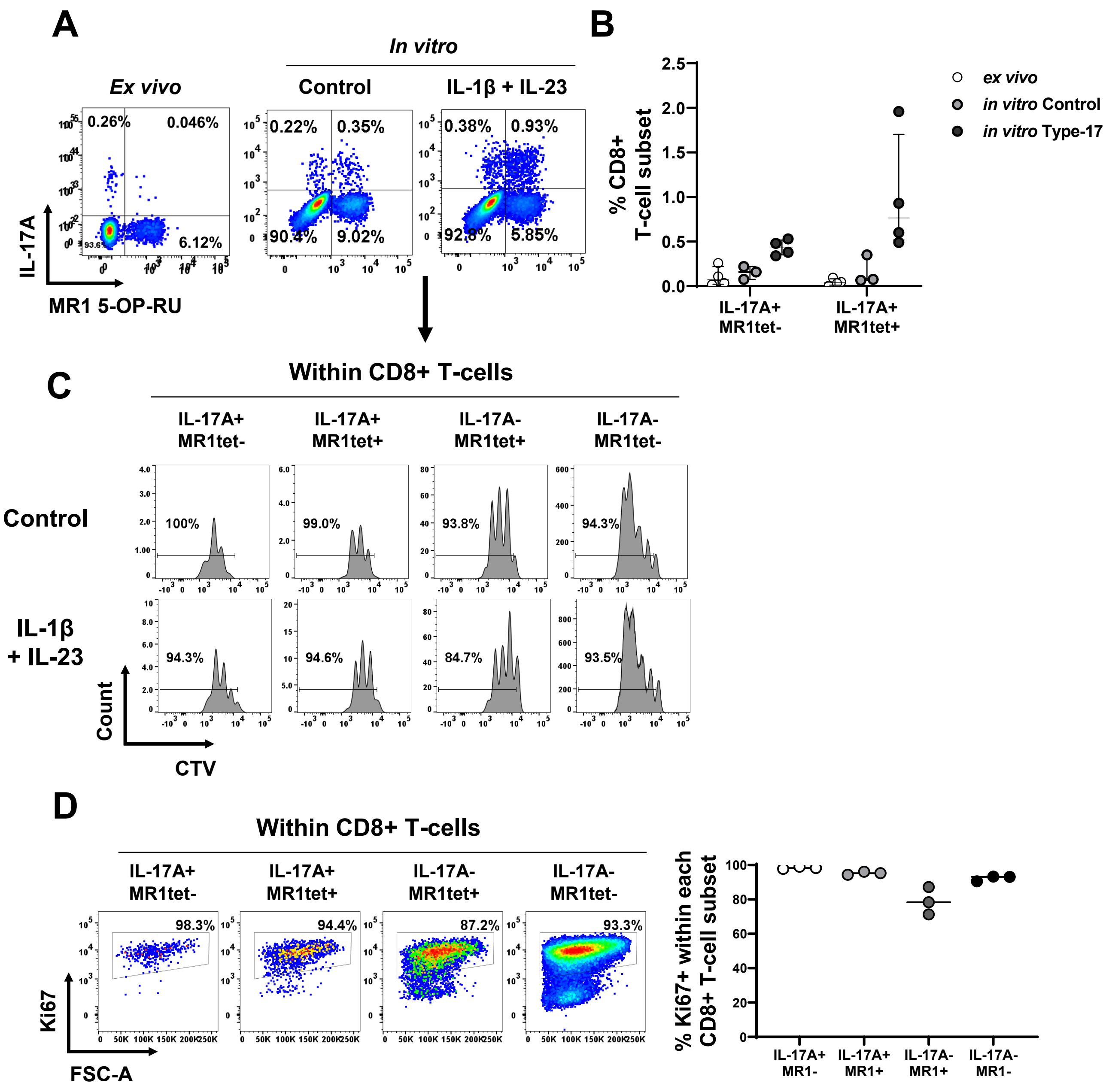


**Supplementary Figure 6. *In vitro*-generated MR1 tetramer- and MR1 tetramer+ IL-17+ CD8+ T cells display comparable proliferative potential.** Healthy donor PBMC were either stimulated *ex vivo* for 3 hours with PMA/ionomycin and GolgiStop or cultured for proliferative assessment. Cells were labelled with CellTrace$^{TM}$ Violet (CTV) then cultured for 3 days with plate-bound anti-CD3 mAb and soluble anti-CD28 mAb in the absence (control) or presence of hrIL-1β and hrIL-23. On day 3, cells were re-stimulated for 3 hours with PMA/ionomycin and GolgiStop and assessed by flow cytometry. **(A)** Representative stainings and **(B)** cumulative frequencies of IL-17A+ MR1 tetramer- and IL-17A+ MR1 tetramer+ cells identified within total live CD8+ T-cells ex vivo or after in vitro culture. **(C)** Histograms depict CTV staining and percentage of proliferating cells after control (upper panel) or type-17 polarising culture (lower panel) in the four indicated populations identified by gating IL-17A vs MR1 tetramer expression within total live CD8+ T-cells (n=1). **(D)** Representative FACS plots and cumulative data (n=3) show intracellular Ki67 staining in each denoted subset within CD8+ T-cells. Data are plotted as median ± IQR.
