## Supplementary Tables1-3. for "Phenotypic, molecular and functional characterisation of human *in vitro*-generated IL-17A+ CD8+ T-cells"

### Supplementary Tables

**Supplementary Table 1**. Details of anti-human monoclonal antibodies and MR1 tetramer used in flow cytometry assessments and T-cell population sorting.

| Target | Supplier | Clone | Fluorochrome conjugates |
|---|---|---|---|
| CD3 | Biolegend | UCHT1 | PE-Cy7 |
| CD4 | Biolegend | SK3 | PerCP-Cy5.5 |
| CD4 | Biolegend | OKT4 | BV421 |
| CD8α | Biolegend | HIT8a | FITC / Pacific Blue |
| CD8α | BD Biosciences | RPA-T8 | BUV395 |
| CD14 | Miltenyi Biotec | REA599 | APC-Vio770 |
| CD19 | Biolegend | H1B19 | APC-Cy7 |
| CD56 | Biolegend | HCD56 | APC |
| CD161 | Biolegend | HP-3G10 | BV421 / BV605 |
| CD161 | Miltenyi Biotec | 191B8 | PE-Vio770 |
| CCR6 | BD Biosciences | 11A9 | APC |
| GM-CSF | Biolegend | BVD2-21C11 | APC |
| Granzyme A | Biolegend | CB9 | PE |
| Granzyme B | Biolegend | GB11 | FITC |
| IL-17A | Biolegend | BL168 | APC / PE |
| IL-17A Cytokine Secretion Assay Detection Kit | Miltenyi Biotec | CZ8-23G1 | APC / PE |
| IL-17F | Miltenyi Biotec | LN2-9C4 | FITC |
| IFN-γ | Biolegend | 4S.B3 | APC / FITC |
| TNF-α | Biolegend | MAb11 | APC / BV605 |
| γδTCR | BioLegend | B1 | APC-Cy7 |
| TCRVα7.2 | Biolegend | 3C10 | APC / BV605 / PE |
| MR1-5-OP-RU or negative control MR1-6-FP tetramers | NIH Tetramer Core Facility | - | PE |

**Supplementary Table 2**. TaqMan primer assay ids selected from ThermoFisher for all 96 genes that were assessed by qPCR array in CSA-FACS sorted *in vitro*-induced IL-17A+ and IL-17A- CD8+ T-cells.

| Gene target | TaqMan assay id | Gene target | TaqMan assay id | Gene target | TaqMan assay id |
|---|---|---|---|---|---|
| **18S** | Hs99999901_s1 | **CCR4** | Hs00747615_s1 | **IL17A** | Hs00174383_m1 |
| **B2M** | Hs99999907_m1 | **CCR5** | Hs00152917_m1 | **IL17F** | Hs00369400_m1 |
| **PPIA** | Hs04194521_s1 | **CCR6** | Hs00171121_m1 | **IL17B** | Hs00975262_m1 |
| **CD4** | Hs01058407_m1 | **CCR7** | Hs01013469_m1 | **IL17C** | Hs00171163_m1 |
| **CD8A** | Hs00233520_m1 | **CCR9** | Hs00246403_m1 | **IL17D** | Hs00370528_m1 |
| **CD8B** | Hs00174762_m1 | **CXCR6** | Hs00174843_m1 | **IL25** | Hs00224471_m1 |
| **KLRB1** | Hs00174469_m1 | **CXCR3** | Hs00171041_m1 | **IL21** | Hs00222327_m1 |
| **DPP4** | Hs00897386_m1 | **CXCR4** | Hs00237052_m1 | **IL22** | Hs01574154_m1 |
| **ICAM1** | Hs00164932_m1 | **ITGAE** | Hs01025372_m1 | **IL26** | Hs00218189_m1 |
| **CD58** | Hs00156385_m1 | **MCAM** | Hs00174838_m1 | **IFNG** | Hs00989291_m1 |
| **CTLA4** | Hs00175480_m1 | **S1PR1** | Hs00173499_m1 | **TNF** | Hs00174128_m1 |
| **ICOS** | Hs00359999_m1 | **RORC** | Hs01076112_m1 | **CSF2** | Hs00929873_m1 |
| **TIGIT** | Hs00545087_m1 | **RORA** | Hs00536545_m1 | **CCL20** | Hs01011368_m1 |
| **PDCD1** | Hs01550088_m1 | **MAF** | Hs00193519_m1 | **GZMA** | Hs00989184_m1 |
| **IL1R1** | Hs00991010_m1 | **IRF4** | Hs01056533_m1 | **GZMB** | Hs00188051_m1 |
| **IL2RA** | Hs00907778_m1 | **STAT3** | Hs01047580_m1 | **PRF1** | Hs00169473_m1 |
| **IL4R** | Hs00166237_m1 | **STAT5A** | Hs00559643_m1 | **IL2** | Hs00174114_m1 |
| **IL6R** | Hs01075667_m1 | **TYK2** | Hs00177464_m1 | **IL4** | Hs00174122_m1 |
| **IL7R** | Hs00902334_m1 | **JAK2** | Hs01078136_m1 | **IL9** | Hs00174125_m1 |
| **IL9R** | Hs00602538_m1 | **AHR** | Hs00169233_m1 | **IL10** | Hs00961622_m1 |
| **IL12RB1** | Hs00538167_m1 | **HIF1A** | Hs00153153_m1 | **IL13** | Hs00174379_m1 |
| **IL12RB2** | Hs01548202_m1 | **FAS** | Hs00163653_m1 | **TGFB1** | Hs00998133_m1 |
| **IL13RA1** | Hs00609817_m1 | **RUNX1** | Hs01021970_m1 | **LTA** | Hs00236874_m1 |
| **IL15RA** | Hs00542604_m1 | **ZBTB16** | Hs00232313_m1 | **ABCB1** | Hs00184500_m1 |
| **IL17RA** | Hs01064648_m1 | **BCL11B** | Hs01102259_m1 | **APOD** | Hs00155794_m1 |
| **IL17RC** | Hs00994305_m1 | **TCF7** | Hs01556515_m1 | **CTSL** | Hs00377632_m1 |
| **IL18R1** | Hs00977691_m1 | **PRDM1** | Hs00153357_m1 | **HOPX** | Hs04188695_m1 |
| **IL21R** | Hs00222310_m1 | **TBX21** | Hs00203436_m1 | **SOCS3** | Hs02330328_s1 |
| **IL23R** | Hs00332759_m1 | **GATA3** | Hs00231122_m1 | **CD7** | Hs00196191_m1 |
| **IL1RL2** | Hs00909276_m1 | **FOXP3** | Hs01085834_m1 | **GPR65** | Hs00269247_s1 |
| **TGFBR2** | Hs00610318_m1 | **EOMES** | Hs00172872_m1 | **ZBTB32** | Hs01004988_m1 |
| **IFNGR1** | Hs00988304_m1 | **ZNF683** | Hs00543184_m1 | **CD5L** | Hs00935901_m1 |

**Supplementary Table 3**. Type-17 signature genes enriched in human *in vitro*-generated IL-17A+ CD8+ T-cells compared with IL-17A- CD8+ T-cells. Differentially expressed genes were identified by paired Student's t-test with Holm-Šídák multiple comparisons with adjusted p<0.05 and having a relative fold change (FC) threshold of ≥1.5 (n=7). Statistically significant genes that did not meet the ≥1.5 criteria but had a FC>1 are denoted in the list above with an asterisk.

| Gene target | IL-17A+ CD8+ T-cells (∆Ct) | | IL-17A- CD8+ T-cells (∆Ct) | | Relative FC (IL-17A+ vs IL-17A-) | Adjusted p-value (p < 0.05) |
|---|---|---|---|---|---|---|
| | Mean | ± SD | Mean | ± SD | | |
| **RORC** | 2.644 | 0.651 | 6.737 | 0.647 | 4.093 | 0.000006 |
| **IL17A** | -2.753 | 0.733 | 6.426 | 0.911 | 9.180 | 0.000008 |
| **IL23R** | 5.130 | 0.925 | 8.561 | 0.925 | 3.431 | 0.000024 |
| **IL17F** | -2.905 | 0.990 | 2.951 | 1.213 | 5.856 | 0.000030 |
| **CCR6** | 7.227 | 1.080 | 11.830 | 1.067 | 4.604 | 0.000065 |
| **RORA** | 0.730 | 0.397 | 2.625 | 0.245 | 1.895 | 0.000156 |
| **MAF** | 3.809 | 0.528 | 6.092 | 0.323 | 2.284 | 0.000176 |
| **IL26** | 2.442 | 0.554 | 6.349 | 0.817 | 3.906 | 0.000328 |
| **MCAM** | 5.105 | 0.713 | 9.377 | 1.029 | 4.272 | 0.001095 |
| **CCL20** | -0.398 | 0.684 | 3.363 | 0.867 | 3.761 | 0.001338 |
| **ICOS** | 1.531 | 0.405 | 3.625 | 0.407 | 2.093 | 0.001726 |
| **ZBTB16** | 3.046 | 0.661 | 6.333 | 0.906 | 3.287 | 0.002619 |
| **CTSL** | 7.391 | 0.484 | 11.135 | 1.104 | 3.744 | 0.002970 |
| **CTLA4** | 0.180 | 0.716 | 3.025 | 0.618 | 2.844 | 0.003765 |
| **IL12RB2** | 2.537 | 0.663 | 4.203 | 0.382 | 1.666 | 0.005018 |
| **IL9** | -0.137 | 1.037 | 3.880 | 1.852 | 4.017 | 0.005454 |
| **IL2RA** | -1.169 | 0.541 | 0.636 | 0.449 | 1.805 | 0.016555 |
| **ICAM1** | 1.950 | 1.000 | 3.882 | 0.892 | 1.931 | 0.019235 |
| **CXCR4** | 2.535 | 0.678 | 4.078 | 0.627 | 1.543 | 0.025260 |
| **IL1R1** | 6.530 | 0.992 | 8.471 | 1.372 | 1.942 | 0.029456 |
| **CXCR6** | 6.575 | 0.677 | 8.630 | 0.680 | 2.055 | 0.031611 |
| **PDCD1** | 4.561 | 0.773 | 6.495 | 0.528 | 1.935 | 0.031627 |
| **GZMB** | 3.228 | 0.614 | 4.365 | 0.923 | 2.335 | 0.039550 |
| **HOPX** | 1.521 | 0.411 | 2.973 | 0.624 | 1.521 | 0.041616 |
| **KLRB1** | -4.332 | 0.944 | -1.997 | 0.408 | 2.267 | 0.044115 |
| **CD58** | 0.421 | 0.635 | 1.942 | 0.323 | 1.137* | 0.037363 |
| **S1PR1** | 3.727 | 0.376 | 5.993 | 0.663 | 1.452* | 0.039550 |
| **IFNGR1** | 3.969 | 0.798 | 5.343 | 0.449 | 1.374* | 0.044392 |
| **ITGAE** | 7.840 | 0.979 | 9.241 | 0.640 | 1.401* | 0.048485 |
